# Clocks and Dominoes: Timing Mechanisms of Embryogenesis

**DOI:** 10.64898/2026.01.25.701537

**Authors:** Yonghyun Song, Brian D. Leahy, Hanspeter Pfister, Dalit Ben-Yosef, Daniel J. Needleman

## Abstract

How developmental timings are regulated is a fundamental open question. Two widely considered mechanisms are the clock, in which an internal timer determines when each stage occurs, and the domino, in which the completion of each stage triggers the next. It is often unclear how to establish either mechanism. Here, we construct a quantitative framework that uses the correlation structure of developmental timings to test the clock and domino mechanisms. We apply this framework to human pre-implantation development by using ~1 million images of 2946 embryos acquired during IVF treatment, establishing mathematical models of developmental rate. We find that a domino mechanism governs the cleavage timings, while a pronuclei fade-triggered clock mechanism governs the morula and blastocyst timings. These results are consistent with the cell cycle oscillator governing the cleavage timings and the accumulation of embryonic gene products or the degradation of maternally deposited factors governing the morula and blastocyst timings. We next investigate the physiological regulators of developmental timing by analyzing how the timings are statistically associated with the clinical pregnancy outcome. While embryos that result in a clinical pregnancy tend to exhibit shorter cleavage timings, this association is primarily driven by patient-specific properties. In contrast, embryo-specific properties independently influence the pregnancy outcome and the cleavage timings, so that factors directly determining implantation potential, such as aneuploidy, can only weakly impact the cleavage timings. Taken together, this work provides a robust framework for decoding developmental timing mechanisms, with significant implications for fundamental biology and clinical practice.

## 1. Introduction

Animals develop from zygote to adulthood through a stereotypic sequence of developmental processes. The regulation of developmental timing is essential for viable growth, as evidenced across diverse taxa. Key examples include the species-specific pace of vertebrate somitogenesis [31], the association between pre-implantation developmental rates and successful pregnancy in mammals [32], and the genetic adaptation of an herbivorous insect to ensure phenological synchrony between embryogenesis and the seasonal emergence of their host plants [47]. However, the principles that govern developmental timing remain unclear. A widely considered model is the developmental clock, in which an internal timer determines when each process occurs [14]. Another prevalent model for biological timing is the domino, in which each process is initiated by the completion of the preceding one, functioning similarly to a checkpoint [34]. It is often unclear how to establish whether the timing of developmental processes is governed by a clock mechanism, a domino mechanism, or a combination of both. Even within the two well-established models of development, the larval molts in nematodes [44] and the midblastula transition in fruit flies [9], the timing mechanisms still remain controversial.

Here, we construct a quantitative framework to analyze the timing of developmental processes, which enables empirical tests of the clock and domino models. Through this framework, we provide a means to elucidate the heterogeneity across individuals and developmental stages. Our work is similar in spirit to studies of cell size control mechanisms, where the *sizer* and *adder* models are tested against measurements of cell size during growth and division [8, 41, 16]. We apply this framework to human pre-implantation development using a dataset of 2946 movies acquired during routine in vitro fertilization (IVF) treatment. In the first 3 days post-fertilization, human embryos undergo three roughly synchronous cleavage cycles to reach the 8-cell stage. As the cells continue to divide, embryos undergo morphological transformations in which the cells tightly adhere to form the morula by the 4^th^ day post-fertilization and subsequently generate the fluid-filled cavity to form the blastocyst by the 5^th^ day post-fertilization [36].

We find that the timings of early cleavages, morula formation, and blastocyst formation cannot be explained by models with a single clock or domino mechanism. Instead, the timings are fully recapitulated by a model in which the cleavage cycles are governed by a domino and the morula and blastocyst formation are governed by a clock that is triggered by pronuclei fade. The regulatory structure of the model highlights that the 4-cell and 8-cell stages are transition points between the early and late developmental programs. Our results also indicate that clocks and dominoes with embryo-specific rates are crucial to explain the timings. We then analyze the factors driving the timing variability and find that pronuclei fade timing is associated with maternal age, which is linked to embryo health stressors. This prompted us to further analyze the association between timing and embryo health, with the clinical pregnancy outcome as the proxy for the latter. Remarkably, we find that patient-specific factors, rather than embryo-specific factors, drive the association between clinical pregnancy outcome and developmental timing. Overall, our approach generates biological and clinical insights into the rate of human pre-implantation development. Moreover, we establish a broadly applicable quantitative framework to study the governing principles of developmental timing.

## 2. Results

### 2.1. Variability of developmental timing in human pre-implantation embryos

To start investigating the timing of pre-implantation development, we used a previously developed image-analysis pipeline to extract the timing of six morphologically defined stages from time-lapse movies of approximately 26,000 human embryos, which were collected during routine treatment cycles from 2012 to 2017 at the IVF unit of Sourasky Medical Center in Tel Aviv, Israel [23](Fig. 1A). Images were captured roughly every 20 minutes across 7 focal planes for up to 5 days of development, yielding up to about 350 images per focal plane for each embryo. The first developmental stage we consider is pronuclei fade, when the envelope of the maternal and paternal pronuclei break down and fade approximately one day post-fertilization (Fig. 1B). The embryos then undergo a series of cell divisions, transitioning through the 2-cell, 4-cell, and 8-cell stages. As the cells continue to divide, the embryo undergoes compaction to reach the morula stage on day 4, and then forms a fluid-filled cavity to reach the blastocyst stage at day 5. During these 5 days of development, human embryos display a wide variety of developmental trajectories, with many arresting before reaching the blastocyst stage [40]. To focus on *standard* developmental trajectories, we analyze embryos that undergo three roughly synchronous cleavage cycles to reach the 8-cell stage and also successfully form the morula and blastocyst (Fig. S1A, S1B). These criteria result in our final dataset of the timing of six developmental stages, from pronuclei fade to blastocyst formation, for 2946 embryos (supplementary section 1).

**Figure 1:**
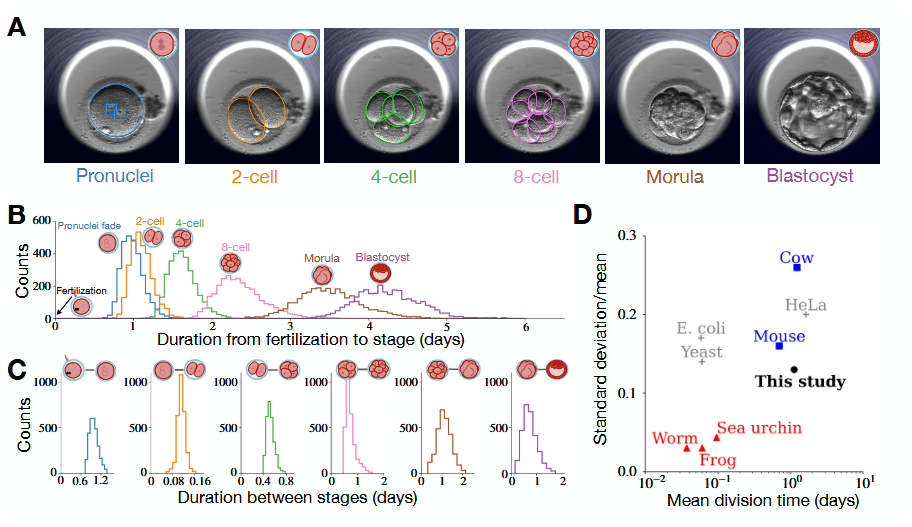
**A** Representative images of a human embryo progressing through the developmental stages of pronuclei fade, 2-cell, 4-cell, 8-cell, morula, and blastocyst. **B** Histograms of the time to each developmental stage from fertilization to blastocyst stage. **C** Histograms of the time intervals between consecutive developmental stages. **D** The mean duration of the first cleavage after fertilization and the associated coefficient of variation (i.e. standard deviation divided by mean) across multiple species. For worm, frog, mouse, cow, and human, the data are based on the duration of the first cleavage after fertilization. For more details, including the references, see Table S1.

Although the order of developmental stages remains consistent across all embryos, the timing of each stage varies significantly (Fig. 1B,C). For example, the blastocyst stage timing ranges from ~3.5 days to ~5.5 days post-fertilization. The relative variability is similar across human preimplantation developmental stages, with the coefficient of variation (standard deviation divided by the mean, CV) of ~0.1, which is consistent with previous measurements [19] (Table S1). To investigate whether human embryos collected for IVF exhibit uniquely large timing variability, we compare the timing of the first cleavage cycle across embryos from different species. The CV of the first cleavage cycle of human embryo (0.13) is smaller than that of cow (0.26), similar to that of mouse (0.16), and larger than that of faster-developing worm (0.04), sea-urchin (0.04) and frog (0.03) (Fig. 1D). Interestingly, the CV of first mammalian cleavage cycles are comparable to that of proliferating unicellular organisms including *Escherichia coli* (0.17), *Saccharomyces cerevisiae* (0.14) and HeLa cells (0.20). Overall, we find that the timing variability in human pre-implantation embryos is comparable to that of other biological systems.

### 2.2. Uniform clock and domino models cannot explain the developmental timings

To explore the mechanisms that govern developmental timing in human embryos, we evaluate two distinct regulatory models: the clock and the domino (Fig. 2A,B).

**Figure 2:**
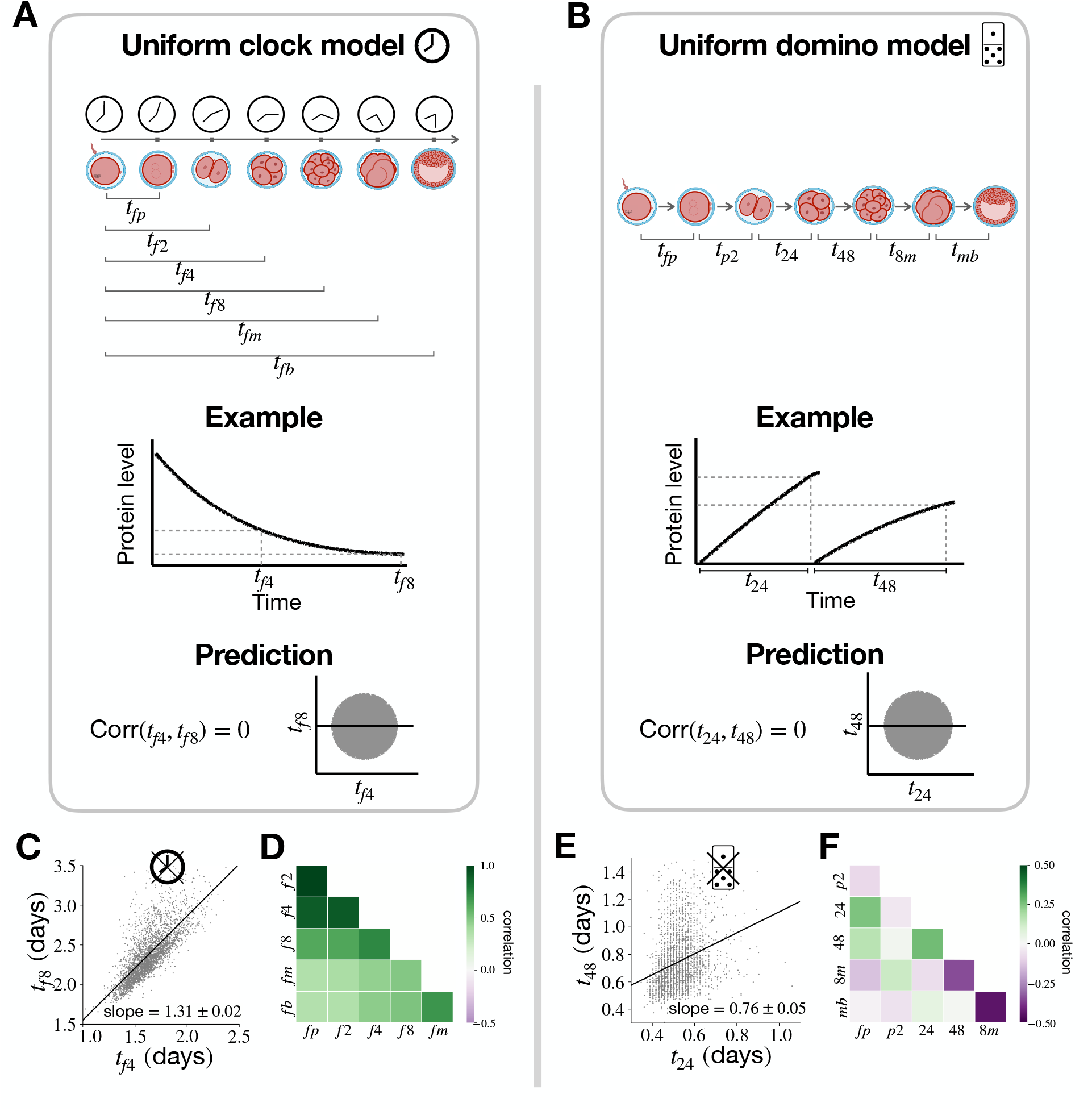
Uniform clock and domino models. **A** Uniform clock model. (Top) Schematic. Stages occur on average at some fixed duration after fertilization. The times to pronuclei fade, 2-cell, 4-cell, 8-cell, morula, and blastocyst from fertilization are respectively denoted by *t*_*fp*_, *t*_*f*2_, *t*_*f*4_, *t*_*f*8_, *t*_*fm*_ and *t*_*fb*_. (Middle) Mechanistic example. *t*_*f*4_ and *t*_*f*8_ are determined by different levels of the maternally deposited protein that degrades over time. (Bottom) Model prediction. Statistical independence between *t*_*f*4_ and *t*_*f*8_. **B** Uniform domino model. (Top) Schematic. Stages occur on average at some fixed duration after the completion of the preceding stage. The duration for fertilization to pronuclei fade, 2 to 4-cell, 4 to 8-cell, 8-cell to morula, morula to blastocyst are respectively denoted by *t*_*fp*_, *t*_*p*2_, *t*_24_, *t*_48_, *t*_8*m*_ and *t*_*mb*_ (Middle) Mechanistic example. The sequential expression of genes whose threshold levels dictate *t*_24_ and *t*_48_. (Bottom) Model prediction. Statistical independence between *t*_24_ and *t*_48_. **C** Plot of *t*_*f*4_ versus *t*_*f*8_. **D** Correlation between all pairs of clock timings. **E** Plot of *t*_24_ versus *t*_48_. **F** Correlation between all pairs of domino timings. In **CE**, *±* denotes the standard error of the slope from linear regression.

In the clock model, a fertilization-triggered time-keeping process dictates the timing of each developmental stage. Since the model assumes that each stage is reached, on average, at a defined duration from fertilization, we designate the timings corresponding to the six stages by *t*_*fp*_, *t*_*f*2_, *t*_*f*4_, *t*_*f*8_, *t*_*fm*_, and *t*_*fb*_ (Fig. 2A top). The clock model could manifest through multiple molecular mechanisms. For example, a maternally deposited protein that undergoes degradation upon fertilization could act as a clock. In this scenario, developmental stages are sequentially triggered when the maternal protein falls below distinct concentration thresholds, resembling a temporal version of the classic French flag model for spatial patterning (Fig. 2A middle)[51].

We first consider the *uniform clock model*, in which the inner clock of each embryo ticks at the same pace in a deterministic manner. However, the clock is read out imperfectly by the embryos, which results in the variability of timing at each stage. Mathematically, we express the timings as

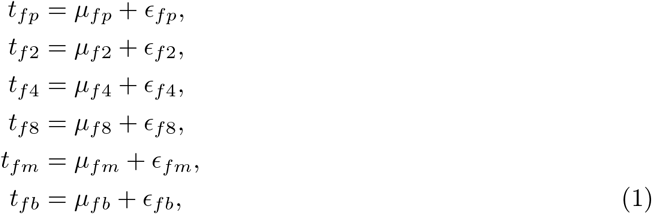

where the *µ* terms are the average values of the respective timings. The *ϵ* terms are the mutually independent random noise, for which no particular distribution is imposed. In the context of the maternally deposited protein example, the *µ* terms embody the threshold concentrations of the degrading maternal protein, while the *ϵ* terms represent the stochastic sensing of protein concentration.

The uniform clock model makes a quantitative prediction that the timings of the different stages must be independent (Fig. 2A bottom). For instance, it predicts a correlation of 0 between the 4-cell and 8-cell stage timings, *t*_*f*4_ and *t*_*f*8_. However, *t*_*f*4_ and *t*_*f*8_ are significantly correlated with Pearson’s correlation coefficient *r* = 0.74 (multiple hypothesis testing adjusted p-value, *p*\* < 10^−10^; Fig. 2C). This correlation was also confirmed with patient-level bootstrapping, and remains significant after accounting for measurement errors (supplementary section 3.3). In fact, all pairs of timings are significantly correlated, which implies that faster embryos tend to reach all the developmental stages more quickly than the slower embryos (*p*\* < 10^−10^; Fig. 2D and Table S3). Therefore, we reject the uniform clock model.

Next, we consider the domino model, where each developmental stage triggers the next sequentially. In this model, each stage occurs on average at a defined duration from the preceding stage, so that the timings are expressed as the following intervals: *t*_*fp*_, *t*_*p*2_[= *t*_*f*2_ *— t*_*fp*_], *t*_24_[= *t*_*f*4_ *— t*_*f*2_],…, and *t*_*mb*_[= *t*_*fb*_ *— t*_*fm*_] (Fig. 2B top). One potential mechanism that may give rise to the domino model is the sequential activation of gene expression (Fig. 2B middle) [27].

We first consider the *uniform domino model*, in which the dynamics for each embryo are on average the same, but each stage occurs with stochastic timing variation. With *µ* and *ϵ* terms defined analogously to the uniform clock model (Eq. 1), we express the timings as

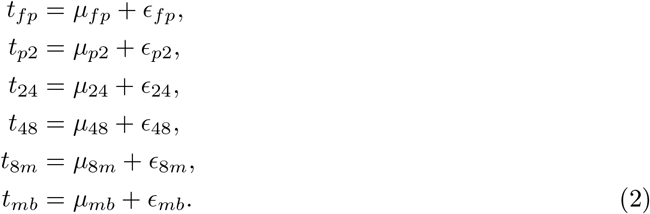

In the sequential gene expression example, the *µ* terms represent the average time to reach the threshold gene expression, while the *ϵ* terms represent the variation in timing due to the stochastic noise of gene expression. The uniform domino model predicts zero correlation between all timings (Fig. 2B bottom). However, the data exhibit significant correlation between multiple pairs, some of which are negatively correlated and others of which are positively correlated (*p*\* < 10^−10^; Fig. 1E,F and Table S3). These results lead us to reject the uniform domino model.

### 2.3. Embryo-specific clock and domino models cannot explain the developmental timings

The results presented in Fig. 2 led us to rule out the uniform clock and domino models, which are perhaps the simplest models of developmental timing. A strong assumption of the uniform models is that all embryos are equivalent, governed by the same underlying dynamical process. However, embryos may have inherently different characteristics that dictate the rate of development, so that some embryos tend to develop faster or slower than others. To address this, we next relax the uniformity assumption and construct models with embryo-specific rates.

We start by considering a clock model in which an embryo-specific latent factor *F*_*c*_ modulates the clock speed. The impact of *F*_*c*_ on the developmental timings is represented in a *causal network diagram*, where directed arrows indicate that *F*_*c*_ affects the timing of each stage (Fig. 3A top). Hence, variations in the rate of the clock cause variations in the timings. In the example with the degrading maternal protein, *F*_*c*_ may correspond to the protein’s characteristic lifetime, *τ*, which varies between embryos (Fig. 3A middle). Embryos with shorter *τ* have shorter timings and embryos with larger *τ* have larger timings.

**Figure 3:**
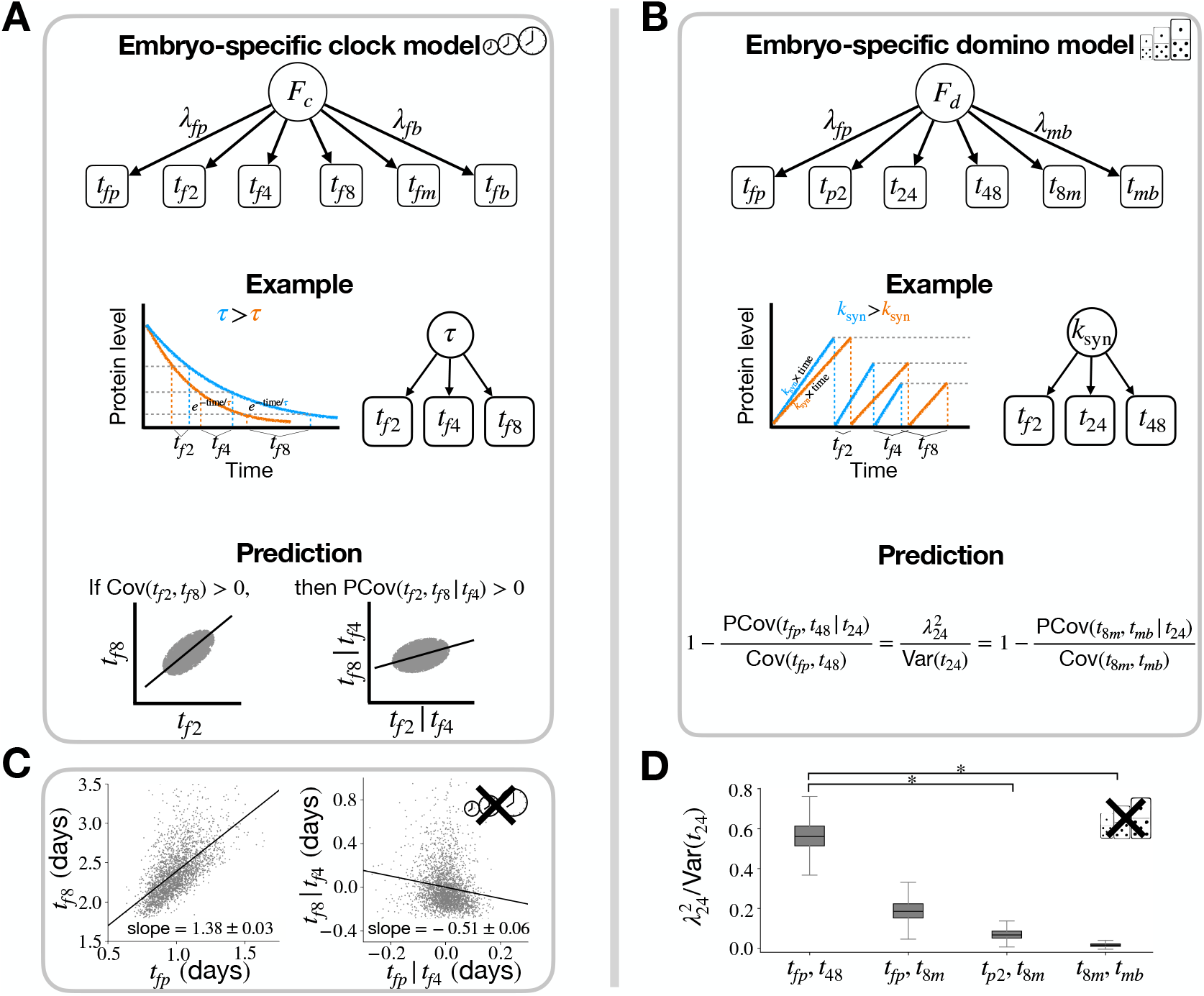
Embryo-specific clock and domino models. **A** Embryo-specific clock model (Top) Schematic. Latent factor *F*_*c*_ influences each clock timing, *t*_*fp*_, *t*_*f*2_, *t*_*f*4_, *t*_*f*8_, *t*_*fm*_ and *t*_*fb*_. (Middle) Mechanistic example. The embryo-specific lifetime of the maternally deposited protein, *τ*, controls the developmental rate. Orange and blue curves represent faster and slower developing embryos, respectively. (Bottom) Model prediction. If two timings, *t*_*f*2_ and *t*_*f*8_ have positive covariance, then the partial covariance after conditioning on a third timing *t*_*f*4_ must also be positive. **B** Embryo-specific domino model. (Top) Schematic. Latent factor *F*_*d*_ influences each domino timing, *t*_*fp*_, *t*_*p*2_, *t*_24_, *t*_48_, *t*_8*m*_, and *t*_*mb*_. (Middle) Mechanistic example. Sequential expression of genes is regulated by *k*_syn_, which governs the rate of gene expression. Blue and orange curves represent faster and slower developing embryos, respectively. (Bottom) Model prediction. The ratio of the partial covariance PCov(*t*_*x*_, *t*_*y*_|*t*_*z*_) and marginal covariance Cov(*t*_*x*_, *t*_*y*_) is independent from the timings *t*_*x*_ and *t*_*y*_. **C** (Left) The covariance between *t*_*f*2_ and *t*_*f*8_. (Right) The partial covariance of *t*_*f*2_ and *t*_*f*8_ after conditioning on *t*_*f*4_. *±* denotes the standard error of the slope. **D** The fraction of variance for *t*_24_ explained by *F*_*d*_ computed with multiple pairs of domino timings. The box denotes the 25% to 75% interquartile range and the whiskers denote 1.5 times the interquartile range. * denotes no overlap between two values from bootstrapping 10^4^ times.

In the *embryo-specific clock model*, we represent the influence of *F*_*c*_ on the timings through linear coefficients *λ*. We assume that *F*_*c*_ is distributed with mean 0 and variance 1 across embryos without loss of generality and express the timings as

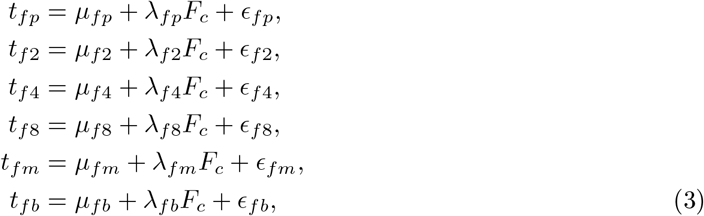

where the *µ* terms denote the population average timings, and the *µ* + *λF*_*c*_ terms embryo-specific average timings. The latent factor, *F*_*c*_, and the noise terms, *ϵ*, are mutually independent random variables, for which we assume no particular distribution.

Previously, we rejected the uniform clock model because, in disagreement with observations, it predicted the timings to be independent of each other (Fig. 2D). In contrast, the systematic variation between embryos in the embryo-specific clock model induced by the latent factor generically leads to correlations between timings. As a result, a Bayesian analysis, in which we assume Gaussian random variables, shows that the embryo-specific clock model is far favored over the uniform clock model with a Bayes factor larger than *e*^100^ (Fig. S2A and Table S2).

We next sought to test additional predictions of the embryo-specific clock model. In the embryo-specific clock model, the correlation between timings are due to the presence of a single latent factor, *F*_*c*_, simultaneously impacting the different developmental stages. Thus, the timings are predicted to be independent when conditioned on *F*_*c*_. While *F*_*c*_ is not directly observed, so it is impossible to condition on *F*_*c*_ in practice, the timings themselves provide imprecise measurements of *F*_*c*_. Conditioning on the timings should therefore partially account for the impact of *F*_*c*_. This intuition can be formalized through the model prediction that the covariance between two timings maintains the same sign when conditioned on a third timing (Eq. S34). For example,

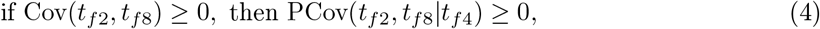

where PCov(*t*_*f*2_, *t*_*f*8_|*t*_*f*4_) is the partial covariance between *t*_*f*2_ and *t*_*f*8_ after conditioning on *t*_*f*4_ (Fig. 3A bottom). This follows because conditioning on *t*_*f*4_ accounts for some, but not all, of *F*_*c*_’s influence on *t*_*f*2_ and *t*_*f*8_. However, the data contradict this. While Cov(*t*_*f*2_, *t*_*f*8_) is positive (*p*\* < 10^−10^), PCov(*t*_*f*2_, *t*_*f*8_|*t*_*f*4_) is negative (*p*\* < 10^−10^). This and other examples in Table S4 rule out the embryo-specific clock model. It is worth noting that the test in Eq. 4 does not require the latent factor or the residual terms to follow any particular distribution, and consistent results are obtained with patient-level bootstrapping. Thus, Eq. 4 provides a general test of the embryo-specific clock model structure.

Next, we consider the *embryo-specific domino model*, in which a latent factor *F*_*d*_, varying from embryo-to-embryo, determines how quickly each developmental process triggers the next (Fig. 3B top). In the sequential gene expression example, *F*_*d*_ could correspond to the overall transcription rate, *k*_syn_. Embryos with smaller *k*_syn_ have longer time intervals between all stages, and embryos with larger *k*_syn_ have shorter time intervals (Fig. 3B middle). Mathematically, we express the timings for the embryo-specific domino model as

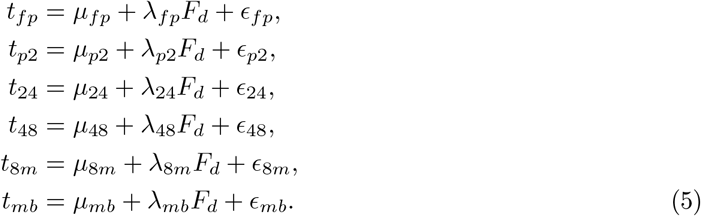

The timing definitions mirror that of the embryo-specific clock model (Eq. 3): the *µ* terms denote the population-average timings, the *λ* terms denote the influence of *F*_*d*_ across developmental stages, and the *ϵ* terms denote the residual variance after accounting for *F*_*d*_.

Unlike the uniform domino model, which was incompatible with the observed correlations between timings, the embryo-specific domino model naturally leads to correlations between timings through the impact of *F*_*d*_. As a result, the embryo-specific domino model is favored over the uniform domino model with a Bayes factor larger than *e*^100^ (Fig. S2B and Table S2).

We further test the embryo-specific domino model by analyzing the effect of *F*_*d*_, which dictates the correlations between all timings. While we cannot directly observe *F*_*d*_, we can partially account for the impact of *F*_*d*_ through the timings themselves. Analogous to Eq. 4, but with clock timings replaced by domino timings in Eq. 5, the embryo-specific domino model predicts that the correlation between two timings maintains the same sign when conditioned on a third timing. Unlike in the embryo-specific clock model, this condition is satisfied for all triples of timings in the embryo-specific domino model (Table S4).

Beyond the preservation of the sign, the model also predicts that the partial covariance is smaller in magnitude than the marginal covariance. The ratio of the partial and marginal covariances quantifies how much variability remains once the latent factor’s influence has been removed. To illustrate this relation more formally, let us take the example of *t*_24_. From Eq. 5, the fraction of *t*_24_’s variance explained by 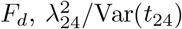, is algebraically equivalent to one minus the ratio of the partial and marginal covariances:

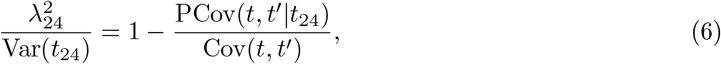

where *t* and *t*′ are distinct timings other than *t*_24_ (Eq. S34). Because the left hand side is determined solely by 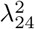 and Var(*t*_24_), it should be the same no matter which pair {*t, t*′} is chosen. In the data, however, the fraction of variance explained for *t*_24_ is ~0.6 when calculated with {*t*_*fp*_, *t*_48_} and ~0 when calculated with {*t*_8*m*_, *t*_*mb*_} (no overlap from bootstrapped intervals, Fig. 3D). Similar inconsistencies occur for *t*_*fp*_, and these conclusions hold after accounting for the measurement errors in the developmental timings (Table S5). These results, which require no assumptions on the particular distribution of *F*_*d*_ and *ϵ* terms, lead us to reject the embryo-specific domino model.

### 2.4. Models that integrate the clock and domino mechanisms explain the developmental timings

The models analyzed thus far assign a single mechanism, either a clock or a domino, to govern the timing of all six developmental stages. However, different stages may be regulated by different mechanisms. We now examine models in which the morula and blastocyst stages follow a timing mechanism distinct from that controlling the earlier cleavage stages.

We start by analyzing the timing of the cleavage stages: pronuclei fade, 2-cell, 4-cell, and 8-cell, but not morula and blastocyst. We consider the embryo-specific clock and domino models of the cleavage timings, which are effectively subsets of the previously discussed embryo-specific clock and domino models (Eqs. 3 and 5). Here, the embryo-specific clock and domino models involve the set of timings {*t*_*fp*_, *t*_*f*2_, *t*_*f*4_, *t*_*f*8_} and {*t*_*fp*_, *t*_*p*2_, *t*_24_, *t*_48_}, respectively, whose correlations are dictated by an embryo-specific latent factor. The embryo-specific clock model of cleavage timings makes an identical prediction to Eq. 4, which requires the sign of the marginal and partial covariances to be equal. However, this is contradicted by the observed values of positive Cov(*t*_*f*2_, *t*_*f*8_) (*p*\* < 10^−10^) and negative PCov(*t*_*f*2_, *t*_*f*8_|*t*_*f*4_) (*p*\* < 10^−10^) (Fig. 3C). Thus, we rule out the embryo-specific clock model for the cleavage stage timings. In contrast, for the embryo-specific domino model of the cleavage timings, the predictions concerning the partial and marginal covariances are fully satisfied by the observed data: the marginal and partial covariances maintain the same sign, and estimates of the fraction of variance explained by the latent factor yield consistent values (entries of Tables S4 and S5 involving only *t*_*fp*_, *t*_*p*2_, *t*_24_ and *t*_48_). For the interval from pronuclei fade to 2-cell stage, *t*_*p*2_, the latent factor-derived variance is negligible (Table S5), which is in accordance with the observation that *t*_*p*2_ is not associated with the other cleavage timings (Table S3). Overall, the cleavage stage timings are well described by the embryo-specific domino model in which a latent factor *F*_*d*_ influences *t*_*fp*_, *t*_24_ and *t*_48_, but not *t*_*p*2_ (Fig. 4A left).

**Figure 4:**
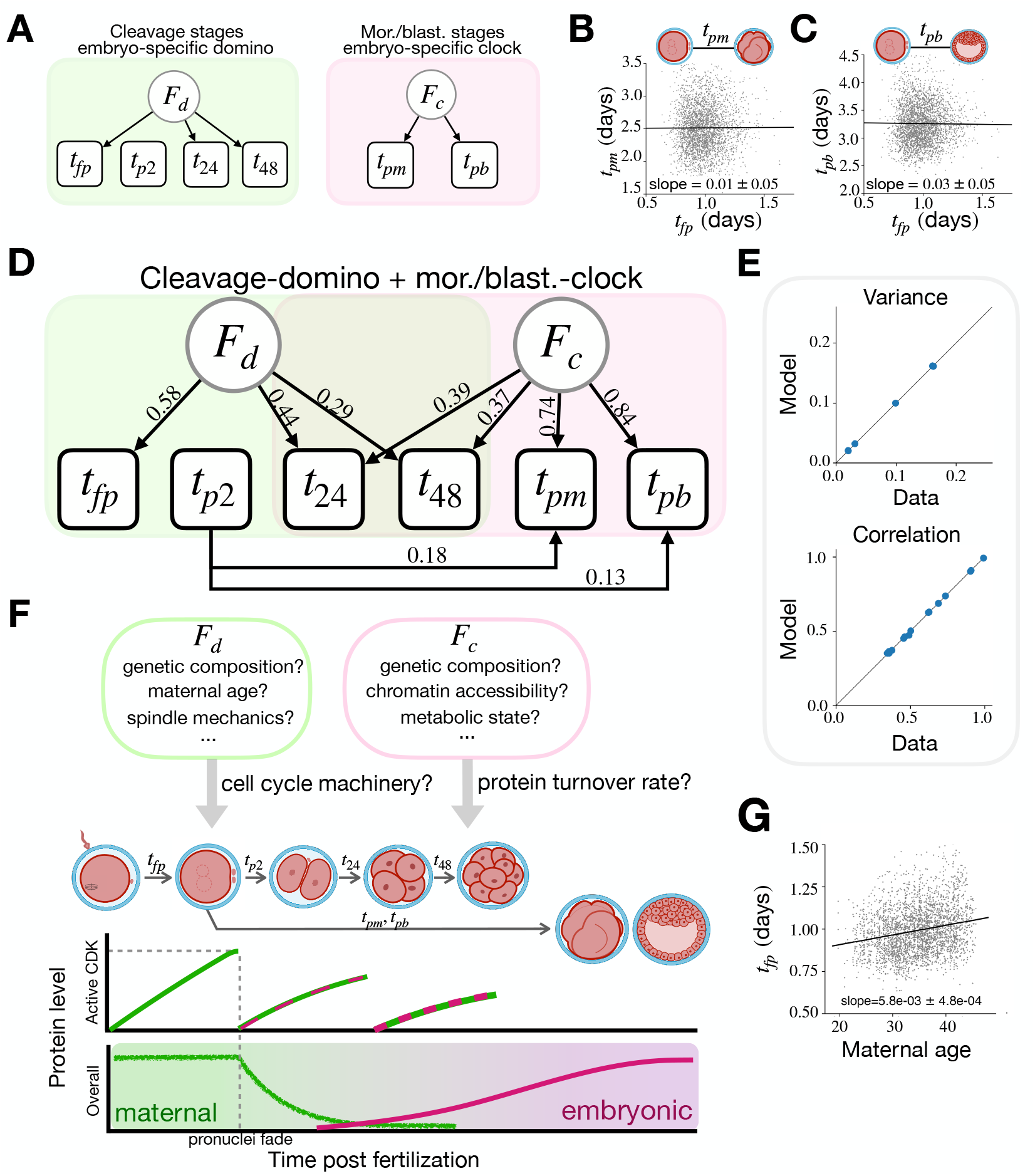
Cleavage-domino + mor./blast.-clock model. **A** Network diagrams of (Left) the domino model for the cleavage stage timings and (Right) the pronuclei fade-triggered clock model for the morula and blastocyst stage timings. **B** Scatter plot of *t*_*fp*_ versus *t*_*pm*_. **C** Scatter plot of *t*_*fp*_ versus *t*_*pb*_. **D** Cleavage-domino + mor./blast.-clock model. The numbers on the arrows are the maximum *a posteriori* probability estimate parameters obtained from fitting the model with normalized timings (supplementary section 3.1). **E** (Top) The variance of timings in the data versus the best-fit model. (Bottom) The correlation of timings in the data versus the best-fit model. **F** Schematic of the possible biological mechanisms that underlie the cleavage-domino and mor./blast.-clock. **G** Plot of maternal age versus *t*_*fp*_. In **BCG**, *±* denotes the standard error of the slope.

Next, we examine morula formation, which has been shown in multiple studies to require the accumulation of embryonic gene products [18, 26, 54, 53]. Assuming that the morula forms at a threshold level of a particular gene product, the simplest model predicts that the interval from the onset of gene expression to morula formation is independent of the timings preceding gene expression. We first test the scenario where the morula-associated gene expression is triggered at the 8-cell stage, which predicts that *t*_8*m*_ is independent from all preceding intervals. However, we observe a significant negative correlation between *t*_48_ and *t*_8*m*_ (*p*\* < 10^−10^; Table S3), which rules out the 8-cell stage as the morula trigger point. In fact, it has been reported that morula formation can occur earlier than the 8-cell stage [30]. Models in which morula formation is triggered at the 4-cell or 2-cell stages are rejected with similar observations (*p*\* < 10^−9^; Table S3). In contrast, the interval from pronuclei fade to morula formation, *t*_*pm*_, shows no association with the preceding timing, *t*_*fp*_ (Fig. 4B). Intuitively, since the variations in timing from fertilization to pronuclei fade are not correlated with the variations in timing from pronuclei fade to morula formation, different factors would underlie those variations. The simplest model consistent with these observations is one in which pronuclei fade triggers the processes dictating morula timing. We obtain similar results for blastocyst timing, with the interval from pronuclei fade to blastocyst formation, *t*_*pb*_, showing no detectable association with *t*_*fp*_ (Fig. 4C). The correlation between *t*_*pm*_ and *t*_*pb*_ is accounted for by an embryo-specific latent factor *F*_*c*_. Overall, these observations are consistent with a regulatory architecture in which the pronuclei fade triggers an embryo-specific clock that regulates the timing of morula and blastocyst formation.

We next sought to construct a single model that combines the two sets of timings discussed above: (i) the cleavage stage timings, {*t*_*fp*_, *t*_*p*2_, *t*_24_, *t*_48_}, governed by an embryo-specific domino, and (ii) the morula and blastocyst stage timings, {*t*_*pm*_, *t*_*pb*_}, governed by an embryo-specific clock. We thus investigate the correlations between the cleavage stage timings and the morula and blastocyst stage timings. Of the four cleavage stage timings, *t*_*fp*_, *t*_24_ and *t*_48_ are associated with each other, while none are associated with *t*_*p*2_. However, *t*_*p*2_ is associated with morula and blastocyst timings (*p*\* < 10^−9^; Table S3). The simplest way to account for this (i.e. adding the fewest number of parameters) is to add directed arrows from *t*_*p*2_ to *t*_*pm*_ and *t*_*pb*_ (Fig. 4D). Of the remaining three cleavage stage variables, *t*_24_ and *t*_48_ are both correlated with *t*_*pm*_ and *t*_*pb*_. The simplest way to account for this is to add arrows from the morula and blastocyst clock latent factor, *F*_*c*_, to *t*_24_ and *t*_48_ (Fig. 4D).

In the resulting *cleavage-domino + mor./blast.-clock model*, the latent factor *F*_*d*_ captures the correlations among the cleavage timings, while *F*_*c*_ links *t*_24_, *t*_48_, *t*_*pm*_ and *t*_*pb*_ (Fig. 4D). The use of two latent factors matches the structure of the data: the correlation matrix of the timings has only two principal components with eigenvalues greater than one, indicating that two factors is enough to capture the bulk of the shared variance (Fig. S3B). Expressed mathematically, the model takes the form

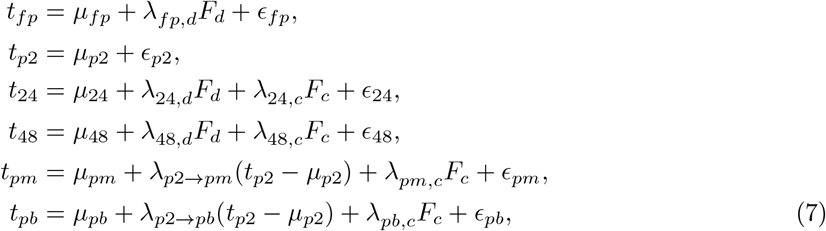

where the *µ* and *ϵ* terms are the average timings and mutually independent noise, respectively. The coefficients *λ*_*fp,d*_, *λ*_24,*d*_ and *λ*_48,*d*_ quantify the influence of the latent factor *F*_*d*_, while *λ*_24,*c*_, *λ*_48,*c*_, *λ*_*pm,c*_ and *λ*_*pb,c*_ quantify the influence of the latent factor *F*_*c*_. The direct influence of *t*_*p*2_ on *t*_*pm*_ and *t*_*pb*_ are denoted by *λ*_*p*2→*pm*_ and *λ*_*p*2→*pb*_, respectively.

When fit to the timings of all the embryos, the cleavage-domino + mor./blast.-clock model reproduces the observed variance and correlations with high fidelity (Fig. 4E). With the best-fit parameters of the model, the influence of *F*_*d*_ decreases for the later cleavages (*λ*_*fp,d*_ *> λ*_24,*d*_ *> λ*_48,*d*_), while the influence of *F*_*c*_ is more prominent for the morula and blastocyst stages than the cleavage stages (*λ*_24,*c*_ ≈ *λ*_48,*c*_ *< λ*_*pm,c*_ ≈ *λ*_*pb,c*_) (Fig. 4D). These observations indicate a transition from the cleavage-domino-(*F*_*d*_) to mor./blast.-clock-(*F*_*c*_) dominated developmental processes, with *t*_24_ and *t*_48_ marking the crossover points.

The cleavage-domino + mor./blast.-clock model architecture paints a physiological picture for the regulation of developmental timing (Fig. 4F). Our analysis indicates that the cleavage stage timings are regulated by a domino mechanism, in line with these timings being determined by cell cycle oscillators that reset with each cell division [10]. In this context, the domino latent factor *F*_*d*_ integrates variables that act on the cell cycle machinery, such as DNA replication rate and spindle mechanics [37, 35]. In contrast, the morula and blastocyst stage timings are regulated by a clock mechanism, which is consistent with regulation by the gradual accumulation of embryonic gene products or the degradation of maternally deposited factors [2]. The associated latent factor *F*_*c*_ could then reflect broadly acting variables such as chromatin accessibility that influences transcription rate [29], and metabolic state that influences translation rate [7, 6]. *F*_*c*_-induced global shifts in the proteome could influence the timing of all events that occur after the clock is triggered at pronuclei fade. This provides an explanation for how *F*_*c*_ also influences the cleavage stage timings *t*_24_ and *t*_48_.

To further study the physiological regulators of developmental timing, we next probe how the developmental timings associate with the patient and treatment cycle variables. We start by considering maternal age, which is consistently documented for each embryo, is well established to influence embryo health, and is associated with aneuploidy [3], metabolic dysfunction [48] and oxidative damage [33]. We find that pronuclei fade time is the only timing that is significantly associated with age, such that an additional decade of maternal age prolongs *t*_*fp*_ by 1.4 hours (Fig. 4G). This age-dependent slowdown suggests that embryo health underlies the variation in pronuclei fade timing. By contrast, two other well documented variables, patient body-mass-index (BMI) and peak oestradiol level during stimulated ovulation, show no significant association with the timings (Table S6).

### 2.5. Patient factors drive the association between clinical pregnancy outcome and cleavage timings

To more directly investigate the link between developmental timing and embryo health, we track the fate of embryos after their transfer to the uterus during IVF treatment (Fig. 5A). We match each embryo’s developmental timing to its clinical pregnancy outcome, defined as the presence of a fetal heartbeat on ultrasound. In line with earlier reports, we find that faster developing embryos are more likely to result in a clinical pregnancy [4, 25, 13, 56]. Specifically, embryos that result in a clinical pregnancy reach the pronuclei fade and complete the 2-to-4-cell interval about 1 hour and 30 minutes faster, respectively, than embryos that do not (cluster-robust linear model, *p*\* = 6.4 × 10^−4^ and *p*\* = 1.7 × 10^−4^; Fig. 5B). The later stage timings are not significantly associated with clinical pregnancy outcome, likely because the sample sizes are not sufficiently large (Table S7). Two broad classes of factors could explain the associations observed for *t*_*fp*_ and *t*_24_. The first involves embryo-specific factors, by which healthier embryos with intrinsically higher developmental competence progress through the cleavage stages more rapidly (Fig. 5C). The second involves factors common to embryos from the same patient or treatment cycle. For example, maternal age or laboratory conditions could prolong the cleavage stages and lower clinical pregnancy rates in tandem.

**Figure 5:**
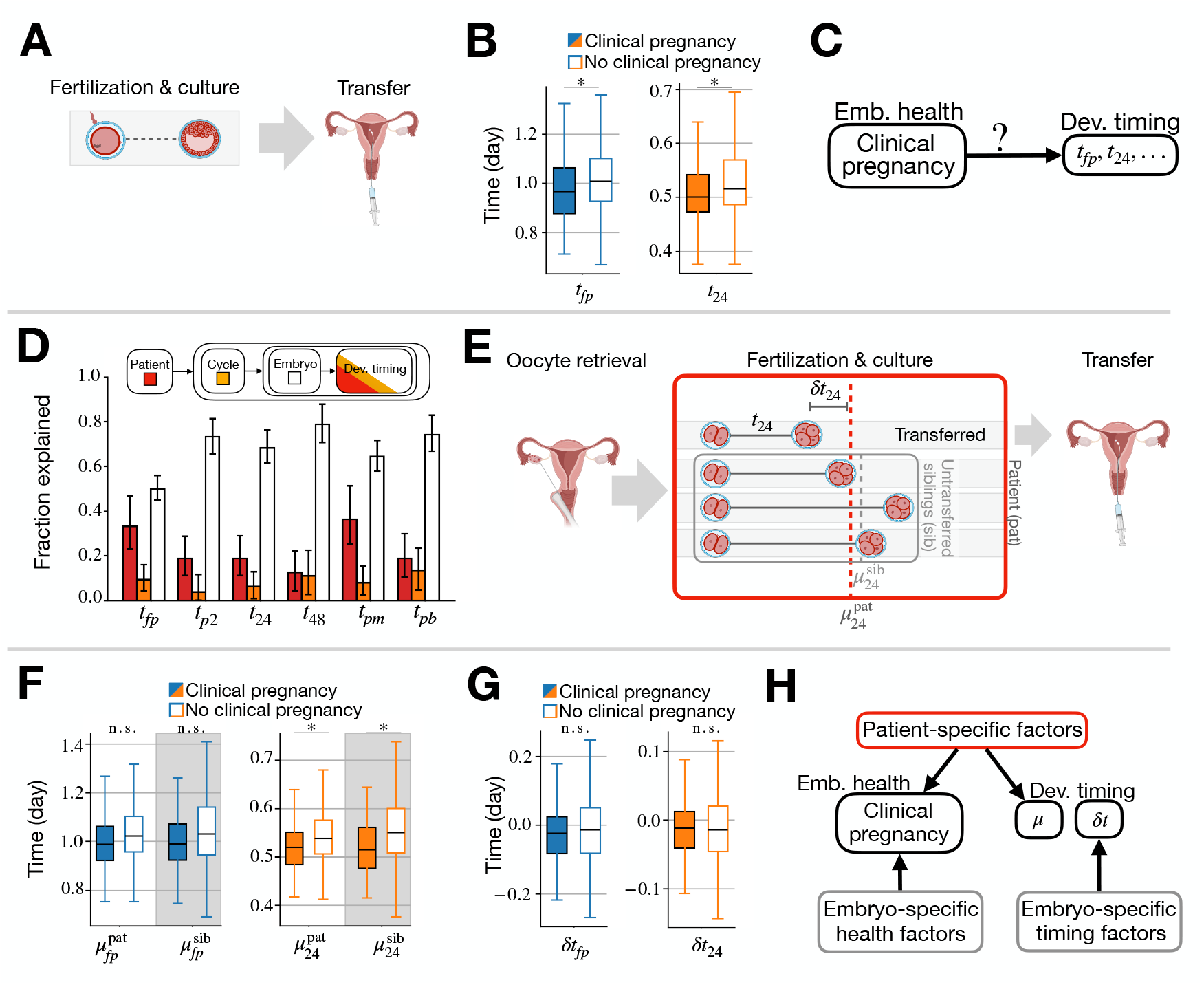
Timing and clinical pregnancy outcome. **A** Simplified schematic for the uterine transfer after culture. **B** Association between the clinical pregnancy outcome and the early cleavage timings, *t*_*fp*_ and *t*_24_. **C** Possible causal diagram for timing and clinical pregnancy outcome. **D** Fraction of variance explained obtained with hierarchical regression. The error bars denote 95% credible intervals. **E** Full schematic for the uterine transfer after culture. Multiple embryos are generated for each cycle, one or more embryos are transferred back to the uterus, and the clinical pregnancy outcome is tested by ultrasound. **F** Association between the clinical pregnancy outcome and the patient-average values. 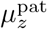 and 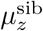 respectively denote the timing of patient-average, and untransferred sibling average, for *z* = *f p*, 24. **G** (Left) Association between the clinical pregnancy outcome and the deviation from patient-average timing. *δt*_*z*_ denotes the deviation of timing from the patient-average for *z* = *f p*, 24. **H** Causal network for timing, clinical pregnancy outcome, patient-specific factors, and embryo-specific factors. In **BFG**, the boxes denote the median and interquartile range, and the whiskers denote 1.5 times the interquartile range.

To assess the impact of patient, treatment cycle, and embryo-specific factors on all the timings, we applied Bayesian hierarchical regression to partition the variance across these three components (supplementary section 3.6). Embryo-specific factors account for half or more of the total variance at every stage (Fig. 5D). Patient factors also explain a substantial portion of the variance, up to 30% in the cases of *t*_*fp*_ and *t*_*pm*_. Lastly, the treatment cycle explains less than 10% of the variability for all stages. Since both embryo-specific factors and patient factors explain a large portion of the timing variability, we next ask which of these factors explains the observed link between clinical pregnancy outcome with *t*_*fp*_ and *t*_24_ (Fig. 5B).

We first analyze the 2-to-4-cell interval, *t*_24_. To separate the patient and embryo-specific factors, we decompose *t*_24_ into

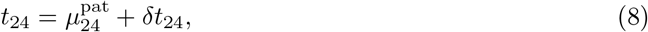

where 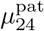 is the average timing of embryos from the same maternal patient, and *δt*_24_ is the deviation of each embryo from 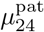 (Fig. 5E). If the observed link between *t*_24_ and clinical pregnancy outcome is driven mainly by patient factors, the clinical pregnancy outcome would be correlated with 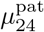 but independent of *δt*_24_. Conversely, if embryo-specific factors dominate, the clinical pregnancy outcome would be correlated with *δt*_24_ but independent of 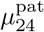.

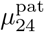 is roughly 30 minutes shorter for embryos that result in clinical pregnancy than for those that do not (*p*\* = 1.9 × 10^−4^; Fig. 5F), which suggests that patient factors contribute to the correlation between *t*_24_ and clinical pregnancy outcome. However, this association may arise simply because 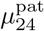 includes the transferred embryos’ *t*_24_. To check this possibility, we consider 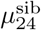, which is the average timing of the sibling embryos from the same patient excluding the transferred embryos. We find that 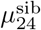 is significantly associated with the clinical pregnancy outcome (*p*\* = 1.6 × 10^−7^; Fig. 5F). In contrast, the within-patient deviation, *δt*_24_, shows no significant difference between embryos that implant and those that do not (Fig. 5G). Thus, patient-specific factors, rather than embryo-specific ones, drive the link between the 2-to-4-cell interval and the clinical pregnancy outcome. Notably, these result manifest even though embryo-specific factors account for a larger share of the timing variability than patient-specific factors (Fig. 5D). The same pattern is observed for pronuclei fade timing *t*_*fp*_: both 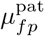 and 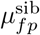 are shorter in embryos that resulted in clinical pregnancy, while the within-patient deviation *δt*_*fp*_ shows no association with clinical pregnancy outcome (Fig. 5F,G). However, the p-values for 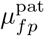 and 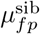 are marginal and do not meet our significance threshold.

In sum, patient-specific factors shared among sibling embryos represent the principal sources of timing variability relevant to clinical pregnancy outcome. This finding ties together earlier reports that linked patient-specific factors either to developmental rate [19, 12] or to clinical pregnancy outcome [15, 46]. By contrast, embryo-specific factors that determine implantation potential exert only a modest influence on the early cleavage timings (Fig. 5H).

## 3. Discussion

In this study, we tested quantitative models of developmental timing against 2946 time-lapse movies of human pre-implantation development. The timings are well explained by a model that combines clock and domino mechanisms, in which a domino mechanism governs the cleavage stage timings, and a pronuclei fade-triggered clock mechanism governs the morula and blastocyst stage timings (Fig. 4). A hand-off between these regulatory modes occurs at the 4-cell and 8-cell stages, which coincide with zygotic genome activation (ZGA) [2]. After establishing this regulatory architecture, we dissected the sources of timing variability and found that although embryo health is associated with differences in timing, it is not causative of these differences (Fig. 5). Patient-specific factors, which are shared among sibling embryos, are the primary drivers of the associations between embryo health and timing. In contrast, embryo-specific properties separately influence embryo health and timing, which suggests that factors directly influencing embryo health, such as aneuploidy and oxidative stress [3, 33], can only weakly influence developmental timing. In the following, we discuss the biological and clinical implications of our findings.

The cleavage-domino + mor./blast.-clock model predicts that the cleavage-domino and the mor./blast.-clock mechanisms can be regulated independently of each other (Fig. 4). This is consistent with observations in mouse, in which manipulating the ploidy or inhibiting DNA replication delays or inhibits the cleavage cycles, but does not affect morula formation and can even accelerate blastocyst formation [5, 43, 20], while in human embryos, a CRISPR knock-out of the gene encoding OCT4 impaired blastocyst formation without affecting the rate of the early cleavage cycles [11].

Our analysis of the developmental timings suggests that pronuclei fade triggers the clock for morula and blastocyst formation (Fig. 4). Since morula and blastocyst formation requires the synthesis of embryonic gene products [18, 11, 53, 57], we hypothesize that the mor./blast.-clock also regulates the timing of ZGA. In human embryos, the major transcriptional burst occurs at the 8-cell stage [2], with low-level transcripts detectable as early as the 1-cell stage [1]. Recent studies suggest that embryonic transcription is initiated through the activity of transcription factors that modulate chromatin accessibility. Some of the candidate ZGA regulators in mouse, such as NR5A2 and OBOX proteins, are already present in the oocyte, and what triggers their activity during embryogenesis remains unclear [21]. Our findings point to processes coincident with pronuclei fade as potential regulators of these transcription factors. For instance, the mitosis-associated chromatin reorganization that follows pronuclei fade may influence the binding of transcription factors to DNA [29, 22]. Alternatively, checkpoint signaling linked to completion of DNA replication and entry into mitosis [37] could trigger maternal mRNA degradation or change global translation rates, thereby affecting the timing of morula and blastocyst formation [42].

Our observations of the association between maternal age and cleavage timings shed light on the mechanisms underlying the cleavage timings. Since age is linked to factors that degrade embryo quality such as aneuploidy [3], metabolic dysfunction [48], and oxidative damage [33], age-dependent changes of the cleavage timings are likely consequences of cell cycle checkpoints being activated by these insults. For example, aneuploidy can prolong cell cycle duration in cells with robust spindle assembly checkpoints [35]. We find that the only developmental stage timing associated with age is pronuclei fade time, *t*_*fp*_, which increases with advanced age (Table S6). We speculate that this may be caused by age-dependent increase in replication stress, which has been shown to delay mitotic entry in human zygotes [37]. The lack of association between age and cleavage timings, *t*_24_ and *t*_48_, indicates that the increased incidence of aneuploidy with age does not result in delays of the cell cycle, consistent with a weak spindle assembly checkpoint. This is consistent with prior experimental studies demonstrating that early mouse embryos have a weak spindle assembly checkpoint [49].

Our findings have implications for designing embryo-selection algorithms. Clinicians aim to transfer only a few embryos at a time to minimize the risks of multiple pregnancies [39, 24], but identifying the most viable embryo remains challenging [17, 38]. When developing embryo-selection algorithms, it may be tempting to correlate embryo-level features with clinical pregnancy outcome and then rank the embryos by the features with the strongest correlations. However, this approach implicitly assumes that each embryo offers an independent observation. Here, we demonstrate that this assumption does not hold, because embryos from the same patient have highly correlated features. While embryos with a shorter 2-to-4-cell interval, *t*_24_, tend to have a higher rate of successful clinical pregnancy, this is a result of certain patients producing cohorts of embryos with both short *t*_24_ and higher clinical pregnancy rates (Fig. 5). Thus, an algorithm based on the shortest *t*_24_ may end up selecting for the best patient rather than the best embryo, providing little help when choosing among sibling embryos from the same patient. Adjusting for commonly considered covariates such as maternal age, BMI, or estradiol level would not eliminate this problem, because *t*_24_ is independent from these variables (Table S6). The patient-specific influence is instead dictated by unmeasured influences, such as genetic or environmental factors. To address these confounders, selection algorithms should evaluate each embryo in the context of its siblings.

Our approach of testing mechanistic timing models through the correlation structure of timings provides a general framework that can be applied to other developmental systems. For classic systems to study timing, such as nematode and fruit fly development, our framework offers a way to evaluate long-standing clock and domino mechanisms [44, 9]. The molecular basis of the clock and domino mechanisms can be further studied by extending this framework to incorporate live-imaging measurements of molecular processes such as gene expression [28] and metabolism [55], and determining their associations and partial associations with the developmental timing variables. Moreover, our framework can be combined with experiments that systematically modulate developmental rate. For example, the rate of mouse pre-implantation development can be controlled with maternal age [52], sex chromosome content [45], and oxygen concentration [50]. Incorporating these variables as additional nodes in the embryo-specific clock and domino models will enable the systematic quantification of their individual and combined influences on developmental rate.

## 4. Materials and methods

Data processing is explained in detail in supplementary section 1, **Description of data**. The full mathematical models are described in supplementary section 2, **Description of models**. The statistical tests employed in the study are further explained in supplementary section 3, **Description of statistical methods**.

## 5. Acknowledgments

We thank all members of the Needleman and Ben-Yosef groups for helpful discussions. Drawings of embryos and the IVF procedure schematic were created at Biorender.com.

## Supplementary Information

### 1 Description of data

#### 1.1 Image source and automated annotation

Time-lapse imaging data were collected using the Embryoscope® system at the Ichilov in vitro fertilization (IVF) center in Tel-Aviv, Israel, as part of routine clinical IVF procedures conducted between 2012 and 2017. Images were acquired in greyscale using Hoffman Modulation Contrast (HMC) microscopy, a form of differential phase contrast microscopy [2]. Images were captured every 20 minutes across 7 focal planes spaced approximately 15 *µ*m apart. Each recording spanned 3 to 5 days, yielding approximately 200 to 400 images per focal plane. Developmental stages, pronuclei fade, 2-cell, 4-cell, 8-cell, morula, and blastocyst formation, were annotated using an automated machine learning pipeline developed by Leahy et al. [4]. Briefly, the pipeline comprises five convolutional neural networks trained to identify morphological features associated with each stage. From these annotations, we extracted the timing of each developmental stage relative to fertilization.

#### 1.2 Selecting embryos with standard developmental trajectories

There were time-lapse images of approximately 26000 embryos collected at the IVF clinic between 2012 and 2017. For this study, we used data from 7972 embryos that were recorded from the zygote stage to the blastocyst stage. The recording may be discontinued before the embryo reaches the blastocyst stage for various reasons, such as embryo transfer or vitrification. From the 7972 eligible embryos, we selected a subset exhibiting what we define as *standard development*. Standard development is characterized by three roughly synchronous cleavage cycles, followed by distinct time points for morula and blastocyst formation. In contrast, *non-standard development* includes irregular features such as tri-polar divisions, cytoplasmic vacuolization, excessive fragmentation, or temporal overlap of key morphological events (e.g., the 8-cell and morula stages occurring simultaneously). The definitions of standard and non-standard development were established specifically for this study, in order to analyze the developmental trajectories that are governed by common underlying biological mechanisms.

In embryos undergoing non-standard development, the time intervals between certain stages are often unusually short or long. For example, the short cleavage stage intervals often correspond to the division of a single blastomere into three or more daughter blastomeres. Thus, to select embryos with standard developmental trajectories, we apply a series of timing-based criteria.

We start with the distribution of the duration between the pronuclei fade and the 2-cell stages, *t*_*p2*_[= *t*_*f2*_ *— t*_*fp*_], which is bimodal (Fig. S1A). We set a lower threshold of *t*_*p2*_ = 0.05 days based on visual inspection of the histogram of *t*_*p2*_ in Fig. S1A. Manual review of 20 randomly selected embryos with *t*_*p2*_ ≤ 0.05 days revealed that 19 of them were mis-annotated by the automated algorithm or the embryo underwent non-standard development. Similarly, an upper cutoff of *t*_*p2*_ = 0.2 days was selected by examining the scatter plot between *t*_*fp*_ and *t*_*f2*_ (Fig. S1B). Here, 15 out of 20 randomly sampled embryos with *t*_*p2*_ *>*= 0.2 days exhibited non-standard development. In contrast, half of the 20 randomly selected embryos with 0.05 *< t*_*p2*_ < 0.2 undergo standard development. A similar bimodal distribution is observed for the time interval between the 2-cell and 3-cell stages, defined as *t*_*23*_[= *t*_*f3*_ − *t*_*f2*_]. Here, we set the lower threshold at *t*_*23*_ = 0.33 (Fig. S1A). Similar duration-based cutoffs are sequentially applied across the following stages: *t*_*p2*_, *t*_*fp*_, *t*_*f2*_, *t*_*23*_, *t*_*f4*_, *t*_*45*_, *t*_*89*_, and *t*_*fb*_. For each cutoff, manual inspection of 20 embryos (or all embryos outside the cutoff when fewer than 20 were available) confirms that more than half show non-standard development (grey group in Fig. S1A). For the final condition based on blastocyst stage timing, only 7 embryos are filtered out, out of which 4 embryos display abnormal morphology. The four most restrictive criteria are highlighted in Fig. S1A. Scatter plots of *t*_*fp*_ versus *t*_*f2*_ and *t*_*24*_ versus *t*_*48*_ in Fig. S1B demonstrate that embryos with standard developmental trajectories form a coherent and clustered group. The full list of duration-based cutoffs, in the order in which they were applied, is as follows:

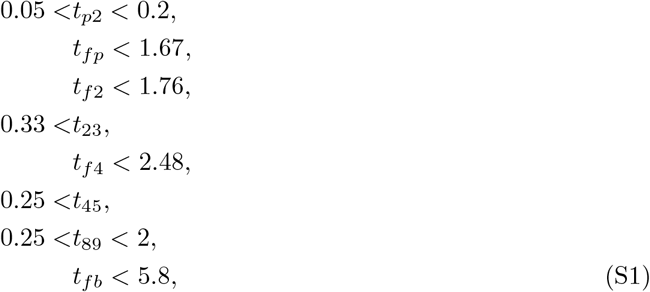

where all the units are days. Overall, all the criteria combine to yield a total of 2946 embryos, which originate from 1178 treatment cycles and 788 patients. 19 out of the 20 embryos sampled from this final group undergo standard development.

**Figure S1:**
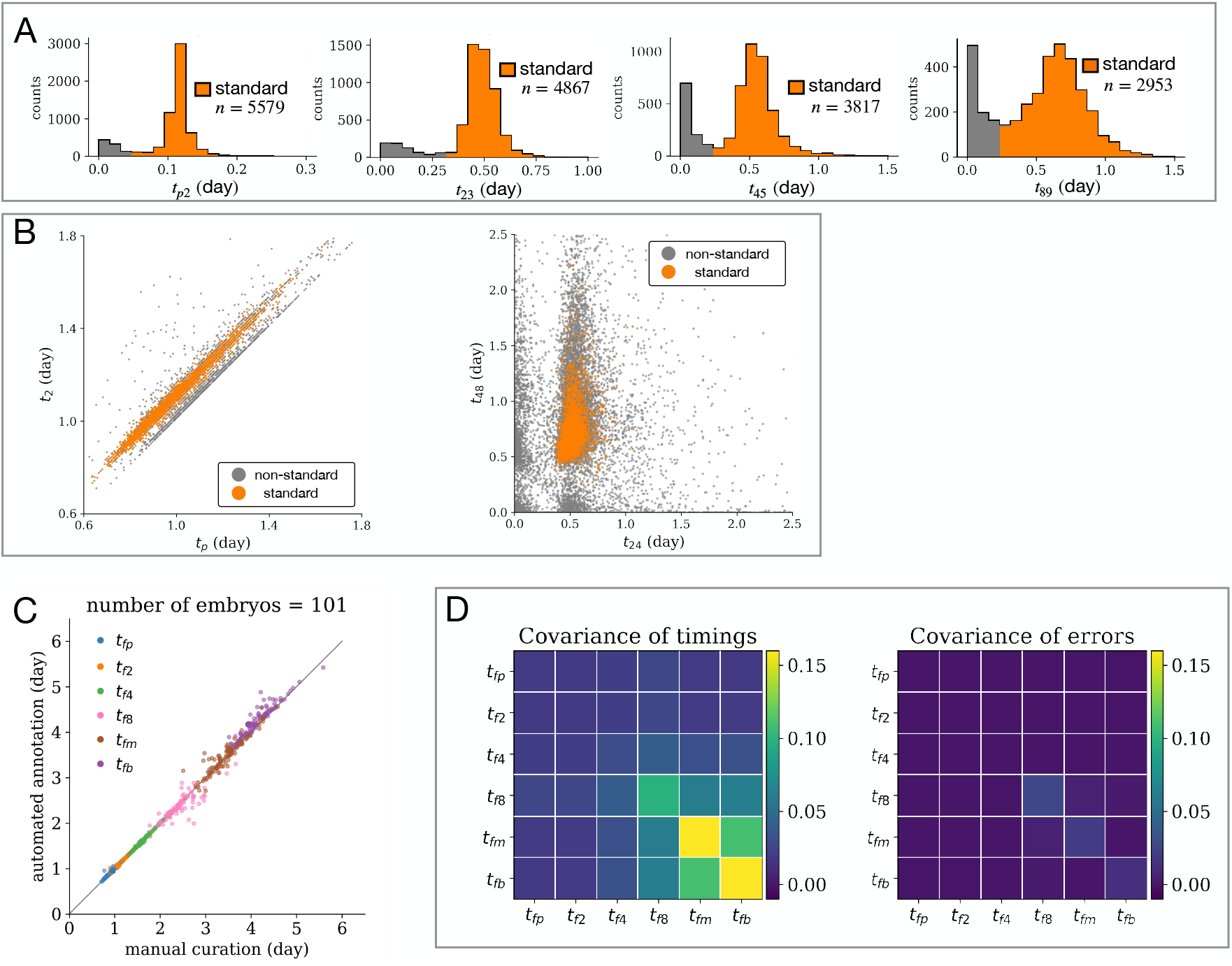
Description of standard developmental trajectories and automated annotation accuracy. **A** Histogram of the time interval between developmental stages, *t*_*p2*_, *t*_*23*_, *t*_*45*_, and *t*_*89*_. Embryos with timings within the orange-shaded region were selected for downstream analysis. **B** Left: Scatter plot of *t*_*fp*_ versus *t*_*f2*_. Right: Scatter plot of *t*_*24*_ versus *t*_*48*_. The embryos selected for analysis are colored in orange, and all others are colored in grey. **C** Comparison between manual annotations by author Y.H.S. and automated stage detection using the machine learning algorithm developed by Leahy et al. [4]. **D** Left: Covariance matrix of developmental stage timings for embryos following standard trajectories. Right: Covariance matrix of annotation errors between the manual and automated methods.

Embryos in the present study are subject to different treatment protocols, which may directly or indirectly influence their developmental timings. For example, the method of fertilization—IVF versus intracytoplasmic sperm injection (ICSI)—has been shown to affect the timing of early cleavage stages [6]. Similarly, pre-implantation genetic diagnosis (PGD) performed at the cleavage stage may directly alter the recorded time to reach the 8-cell stage by altering the number of blastomeres. However, we do not directly model these treatment-specific effects explicitly, as the decision to apply a particular treatment is often influenced by a complex set of factors related to the patient and embryo.

For the hierarchical regression analysis of the influence of patient factors on developmental timings and for the association study between timings and clinical pregnancy outcome, different subsets of embryos were used. For further details on embryo inclusion criteria, statistical methods, and results, see the respective supplementary sections.

#### 1.3 Association between timings and clinical pregnancy outcome

We provide details on the inclusion criteria for evaluating the association between developmental timings and the clinical pregnancy outcome after transfer.

The clinical pregnancy outcome is quantified from the number of heartbeats recorded from the ultrasound examination. We augment and adjust the heartbeat counts by using other clinical records as follows. (i) If the heartbeat count was recorded and larger than the number of embryos transferred, then the heartbeat count is changed to the number of embryos transferred. This is so that we ignore multiple gestation events from the same embryo. (ii) If the heartbeat count was missing and the beta hCG measurement was negative (i.e. less than 25 pg per mL), then the heartbeat count is set to 0. Similarly, in cases where the clinician reported only a chemical pregnancy for the treatment cycle without a fetal heartbeat detection, the heartbeat count is set to 0. (iii) If the number of children born from the treatment cycle was greater than or equal to the number of transferred embryos, the heartbeat count is set to the number of transferred embryos. These adjustments increased the number of usable clinical pregnancy outcome measurements whenever other clinical records provided unambiguous evidence.

To maximize the statistical power for this analysis, we include embryos regardless of whether they were imaged throughout the full duration of pre-implantation development. Each embryo must still follow the standard developmental trajectory criteria described in supplementary section 1.1, but only for the stages captured in the movie. For example, consider embryo *i*, whose video recording spans from the 2-cell stage to the 4-cell stage and is subsequently transferred on day 2 post-fertilization. We include embryo *i* in the analysis only if it satisfies all relevant criteria that apply to the recorded stages. Specifically, if 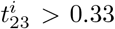 days, 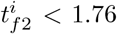 days, and 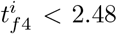 days (Eq. S1). We do not apply any criteria involving developmental stages outside the observed window.

Next, we limit the dataset to treatment cycles whose clinical pregnancy outcomes are unambiguous. After single embryo transfers, one fetal heartbeat signifies success and none signifies failure. After transfers of two embryos, we retain only cycles with either zero heartbeats or two distinct heartbeats, because these correspond to neither or both embryos leading to clinical pregnancy, respectively. Cycles with exactly one heartbeat are excluded since the individual fates of the embryos cannot be assigned with certainty. Analogous criteria are applied for the three and four transfer treatment cycles. Consequently, every treatment cycle included in the analysis offers a clear all-or-none clinical pregnancy outcome for the transferred embryos.

Finally, when computing the association between clinical pregnancy outcome and patient-average timings, we retain only those patients with at least two transferred embryos with standard developmental trajectories. When computing the mean timing of non-transferred siblings, we retain only those patients with at least one transferred embryo and at least one non-transferred sibling, each with standard developmental trajectories. The total number of embryos that satisfy all of these conditions for each analysis is reported in Table S7.

#### 1.4 Errors associated with the automated annotations

Although the automated annotation algorithm developed by Leahy et al. has been shown to be highly accurate, occasional mis-annotations still occur [4]. A systematic bias from the automated annotations may affect our conclusions regarding correlations among the timings. Consider the correlation between the two timings *t* and *t*′. With corresponding annotation errors *ϵ* and *ϵ*′, the measurements of *t* and *t*′ can be expressed, respectively, as 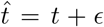 and 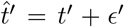. Then, the relationship between the true and measured variance and covariance can be written as follows:

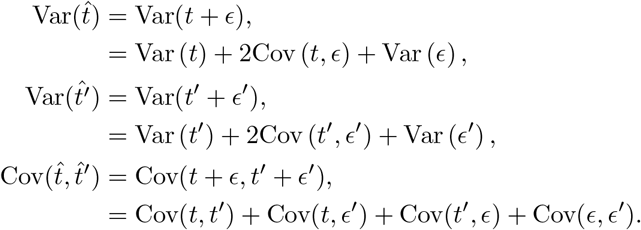

We assume that the errors of the automated annotations are independent from the true duration since they respectively represent the unrelated properties of image-processing and developmental dynamics. Then, Cov(*t, ϵ*) = Cov(*t*′, *ϵ*′) = Cov(*t, ϵ*′) = Cov(*t*′, *ϵ*) = 0, which allows the true correlation to be inferred from the measured correlation by

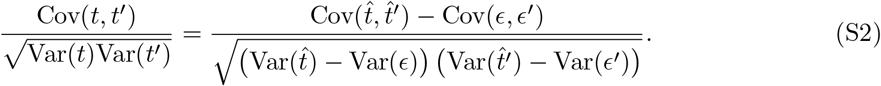

To quantify the variance and covariance of annotation errors, 101 embryos were randomly selected from the 2,946 embryos exhibiting standard development. The developmental stages of these embryos were manually annotated by author Y.H.S. The measurement error for each timing variable is defined as the difference between the manually annotated value and its corresponding automated annotation (Fig. S1D). We find that both the variance and covariance of these measurement errors are substantially smaller in magnitude compared to the overall variability observed across embryos, indicating that automated annotation introduces minimal noise relative to the biological variation (Fig. S1E).

In the analysis of correlations by bootstrapping, we use Eq. S2, and find that the measurement error does not affect the conclusions in the main text regarding the significance of correlations between the timings. For more detail, see the supplementary sections 3.3 and 3.4.

**Table S1:** Extended statistics of cell cycle durations in minutes, corresponding to Fig. 1D in the main text.

| Organism | Mean | Stan. Dev. | Coef. Var. | Reference |
| --- | --- | --- | --- | --- |
| *Escherichia coli* | 84.5 | 14.4 ($n = 4558$) | 0.17 | [14] Fig. S5A |
| *Bacillus subtilis* | 75 | 20.3 ($n = 3637$) | 0.27 | [14] Fig. 2A |
| *Saccharomyces cerevisiae* | 87 | 12 ($n = 116$) | 0.14 | [12] Table S7,8 |
| HeLa S3 | 2359 | 402 ($n > 100$) | 0.20 | [11] Fig. 3A |
| *Caenorhabditis elegans* embryo C | 53.3 | 1.8 ($n = 20$) | 0.03 | [1] Table S1 |
| *Hemicentrotus pulcherrimus* embryo | 138 | 6 ($n > 600$) | 0.04 | [9] Table 1 |
| *Xenopus laevis* embryo | 86.5 | 2.7 ($n = 88$) | 0.03 | [13] Table S1 |
| Mouse embryo ** | 1008 | 162 ($n = 183$) | 0.16 | [5] Table 1 |
| Bovine embryo $t_{f2}$ † | 1770 | 454 ($n = 431$) | 0.26 | [15] Fig. 2B |
| Human embryo $t_{f2}$ no live birth †† | 1572 | 158 ($n = 159$) | 0.10 | [3] Fig. 1 |
| Human embryo $t_{fp}$ | 1612 | 212 ($n = 2946$) | 0.14 | this study |
| Human embryo $t_{f2}$ | 1612 | 212 ($n = 2946$) | 0.13 | this study |
| Human embryo $t_{f4}$ | 1612 | 212 ($n = 2946$) | 0.11 | this study |
| Human embryo $t_{f8}$ | 1612 | 212 ($n = 2946$) | 0.13 | this study |
| Human embryo $t_{fm}$ | 1612 | 212 ($n = 2946$) | 0.11 | this study |
| Human embryo $t_{fb}$ | 1612 | 212 ($n = 2946$) | 0.09 | this study |

\*\* Fresh embryos fertilized by ICSI.
† Fresh embryos fertilized by ICSI. The reported median is taken as the mean value. Standard deviation is estimated by transforming the 25% to 75% quartile range into 1.35 standard deviations.
†† Reported median is taken as the mean value. Standard deviation is estimated by transforming the range between the whiskers in the box plot into 5.4 standard deviations. Both ICSI and IVF fertilization methods were used.
For † and ††, WebPlotDigitizer [10] was used to infer the numerical values from the plots.

### 2 Description of models

#### 2.1 Mathematical form of the causal network models

We provide the mathematical details related to all the models in the main text. In the uniform clock model, the times to the six key morphological events are denoted by the vector 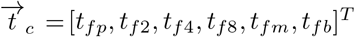. In all model-fitting procedures, we normalize the timing variables by subtracting the respective mean and dividing by the respective standard deviation. The normalized variables are denoted with the tilde sign, so that 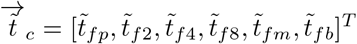 is the normalized timing vector with mean 0 and variance 1 for all its elements. The individual timings are expressed as

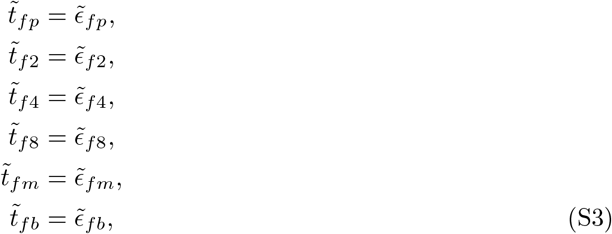

where the residuals 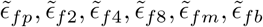 are independently distributed with mean 0 and variance 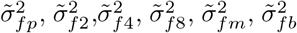. Let 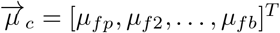 be the mean vector of the non-normalized timings, and Ψ_*c*_ the diagonal matrix composed of the variance of the non-normalized timings. Note that we do not generally require the residuals to follow a particular distribution.

To compute the likelihoods associated with the model, we assume that the residuals follow the Gaussian distribution. Then, the non-normalized timing vector, 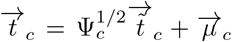, follows the multivariate Gaussian with mean 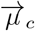 and covariance

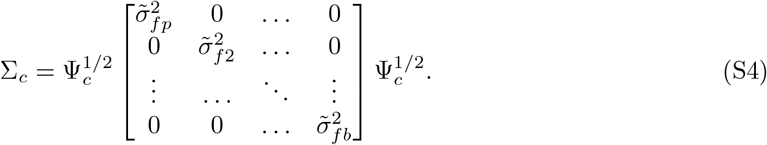

The non-normalized and normalized timing vectors for the uniform domino model are denoted by 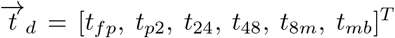 and 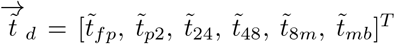, respectively. The normalized timings are expressed as

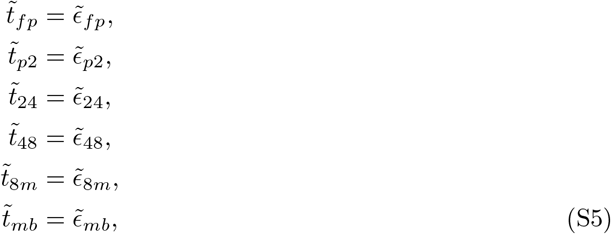

where the residuals 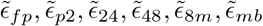 are independently distributed with mean 0 and variance 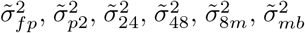. Similarly to the clock model, we do not necessarily specify the residuals to follow a particular distribution. The domino and clock timings are related to each other by 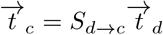, where the matrix *S*_*d*→*c*_ is given by

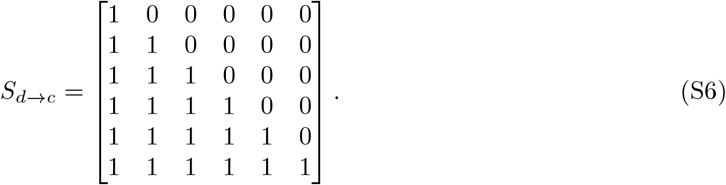

If the residual terms follow the Gaussian distribution, the clock timings 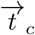 expressed by the domino timings 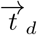 follow the multivariate Gaussian with mean

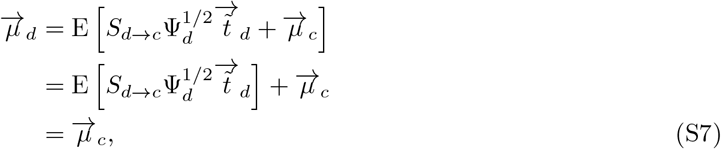

and covariance

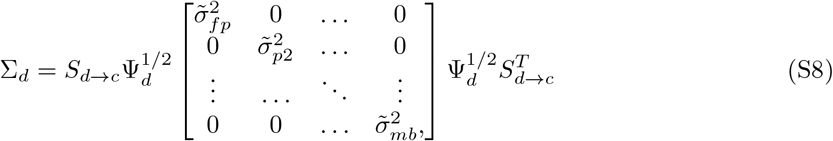

where Ψ_*d*_ is the diagonal matrix composed of the variance of the non-normalized domino timings. Here, the mean timings derived from the domino model, 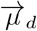, is by definition identical to that of the clock model, 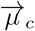. However, the covariance matrices are different between the clock and domino models. ∑_*c*_ is a diagonal matrix, but ∑_*d*_ has non-zero off-diagonal elements because of *S*_*d*→*c*_.

For the embryo-specific clock model, the vector of linear coefficients for the six timings are denoted by 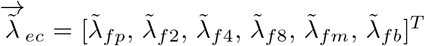. Again, ~ signifies that 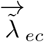 is defined for the normalized timings. Assuming that the embryo-specific latent factor *F*_*c*_ is distributed with mean 0 and variance 1, the normalized timings are expressed as

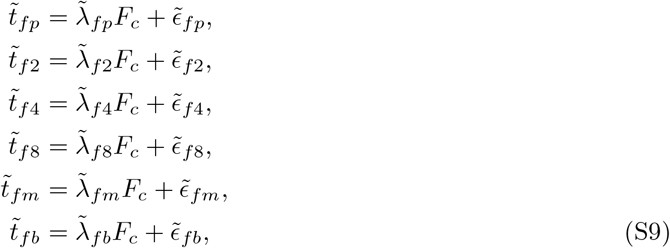

where the 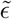 terms are the mutually independent noise terms remaining after conditioning on *F*_*c*_ (Eq. S3). Given *F*_*c*_ = *f*_*c*_, the expected clock timings are expressed as

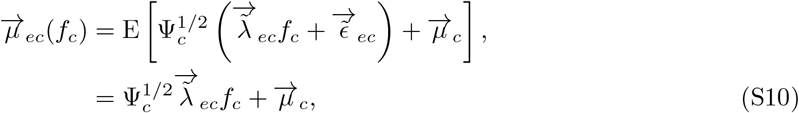

where 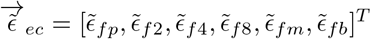 and Ψ_*c*_ is the diagonal matrix composed of the variance of the non-normalized clock timings. Assuming that the residual terms follow the Gaussian distribution, the clock timings given *F*_*c*_ = *f*_*c*_ follows the multivariate Gaussian distribution with mean 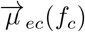 and covariance

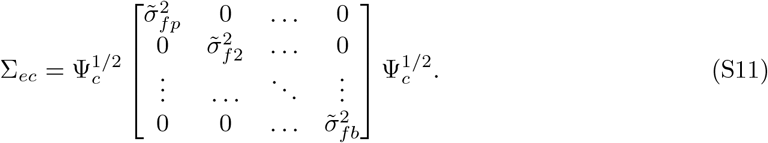

For the embryo-specific domino model, the normalized timings are expressed as

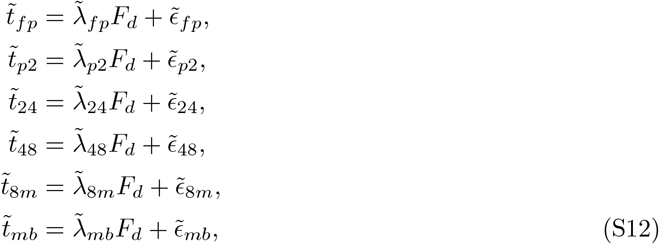

where *F*_*d*_ denotes the domino latent factor distributed with mean 0 and variance 1, the 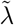 terms denote the linear coefficients for the latent factor *F*_*d*_, and the *ϵ* terms denote the residual noise after conditioning on *F*_*d*_. Then, the expected clock timings given *F*_*d*_ = *f*_*d*_ can be expressed as

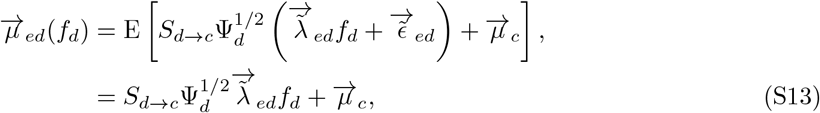

where 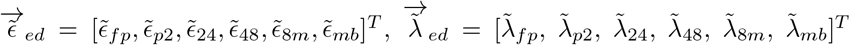 and Ψ_*d*_ is the diagonal matrix composed of the variance of the non-normalized domino timings. With the residual terms following the Gaussian distribution, the domino timings given *F*_*d*_ = *f*_*d*_ follow the multivariate Gaussian distribution with mean 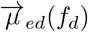 and covariance

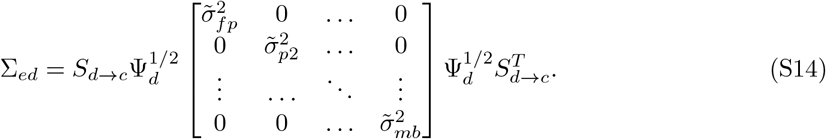

The cleavage-domino + mor./blast.-clock model involves the timing vector denoted by 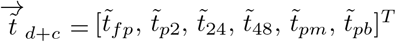, where each of the normalized timings are expressed as

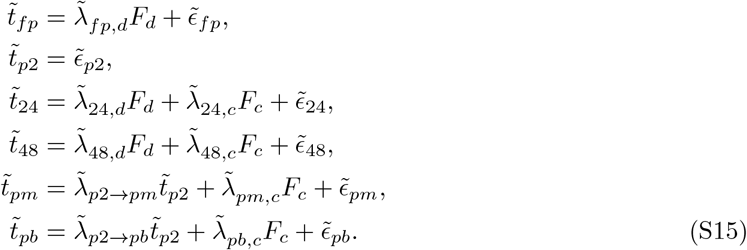

*F*_*d*_ and *F*_*c*_ are latent factors with mean 0 and variance 1, and the 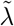 terms are the linear coefficients, so that the vector of linear coefficients for *F*_*d*_ and *F*_*c*_ can be expressed as 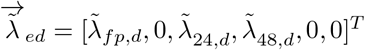 and 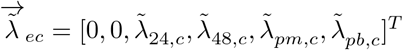, respectively.

Given the latent factors *F*_*d*_ = *f*_*d*_ and *F*_*c*_ = *f*_*c*_, the expected clock timings are expressed as

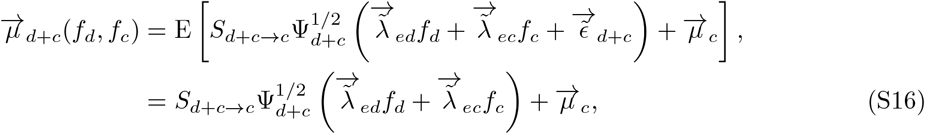

where Ψ_*d+c*_ is the diagonal matrix of the variance for the non-normalized model timings, and

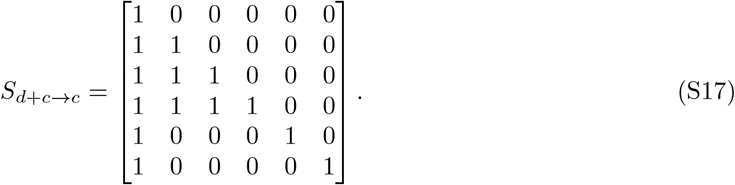

Assuming that the residuals follow the Gaussian distribution, the clock timings follow the multi-variate Gaussian with mean 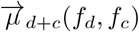 and covariance

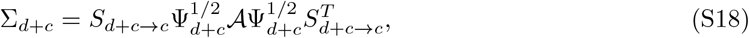

where

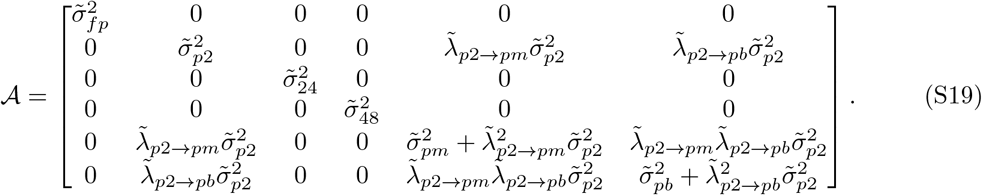

### 3 Description of statistical methods

#### 3.1 Model likelihood

We fit the models by finding the parameters that maximize the Bayesian posterior given the data. The likelihood is evaluated with the non-normalized clock timings, even though the models are expressed in terms of the timings normalized by their respective mean and standard deviation. This way, the linear coefficients provide a means to interpret the relative magnitude of the impact of latent factors on the timings. To evaluate the likelihoods, the latent factors are assumed to follow the standard normal distribution, and the residuals are also assumed to be Gaussian distributed. The prior for the linear correlation coefficients are specified by a standard normal distribution, and the prior for the residual variance is specified by a log-normal distribution.

For the uniform clock model, the fitted parameters are the variance of the residuals, which are collectively denoted by the set 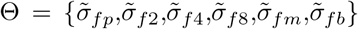. The posterior probability of the parameters given the data **T** from the set of embryos *E* is

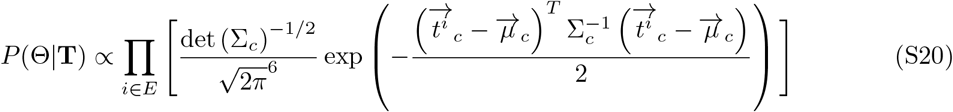

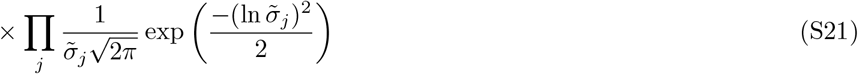

where *i* runs through the embryos and *j* runs through the six different clock timings. The first term (Eq. S20) corresponds to the likelihood integrated over the set of embryos *E*, and the second term (Eq. S21) corresponds to the prior of the variance. 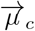 is the mean of the non-normalized clock timings, and the covariance matrix ∑_*c*_ is defined in Eq. S4.

For the uniform domino model, the posterior probability of the parameters, 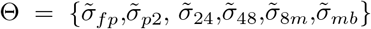, is proportional to

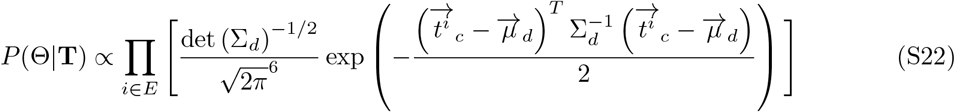

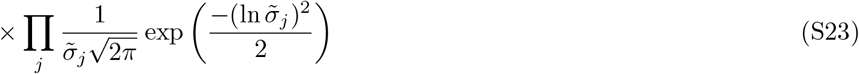

where *i* runs through the embryos, *j* runs through the six domino timings, 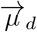 is as defined in Eq. S7, and ∑_*d*_ is as defined in Eq. S8. The first term (Eq. S22) corresponds to the likelihood integrated over the set of embryos *E*, and the second term (Eq. S23) corresponds to the prior of the standard deviations.

For the embryo-specific clock model (Eq. S9), the posterior probability of the parameters 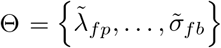 is

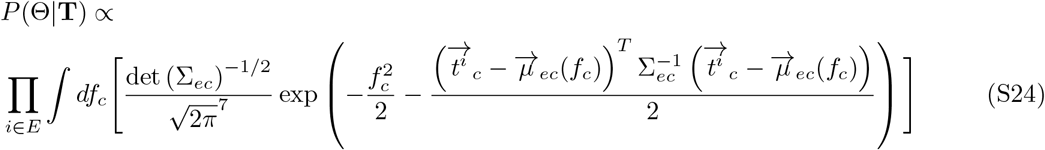

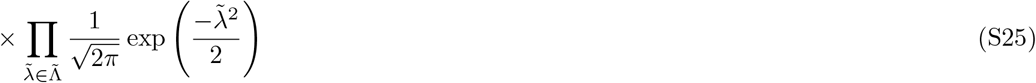

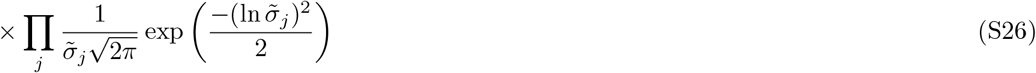

where 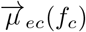 is as given in Eq. S10, ∑_*ec*_ is given in Eq. S11, 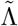 denotes the set of all linear coefficients, and *j* runs through the six different clock timings. The first term (Eq. S24) corresponds to the likelihood integrated over the latent factor *F*_*c*_ and embryos *E*, the second term (Eq. S25) the prior of the linear coefficients, and the third term (Eq. S26) the prior of the standard deviations.

The embryo-specific domino model (Eq. S12) is evaluated analogously to the embryo-specific clock model, using the following expression for the posterior,

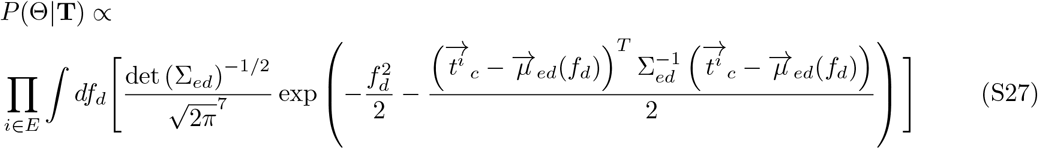

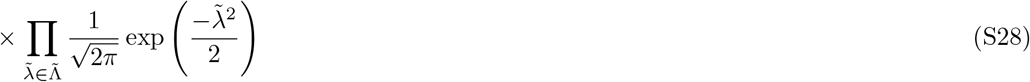

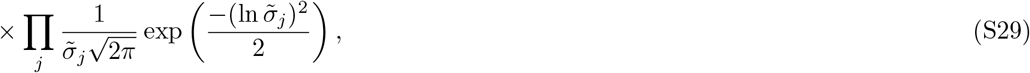

where 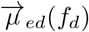 is given in Eq. S13, and ∑_*ed*_ is given in Eq. S14. The first term (Eq. S27) corresponds to the likelihood integrated over the latent factor *F*_*d*_ and embryos *E*, the second term (Eq. S28) the prior of the linear coefficients, and the third term (Eq. S29) the prior of the standard deviations.

The posterior of the cleavage-domino + mor./blast.-clock model (Eq. S15) is expressed as

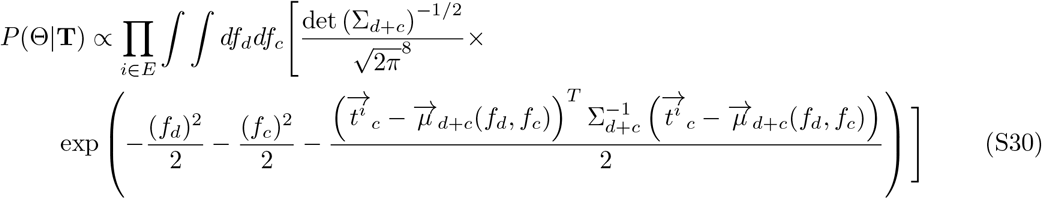

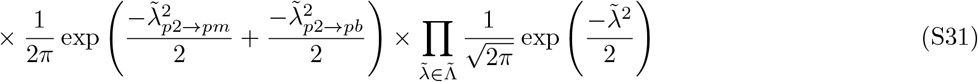

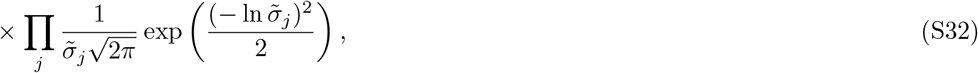

where the first term (Eq. S30) integrates over the two latent factors, *F*_*d*_ and *F*_*c*_. The expressions for 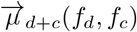 and ∑_*d+c*_ are found in Eq. S16 and Eq. S18, respectively. 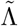 denotes the set of all linear coefficients from latent factors to the timings and *j* runs through all the developmental stages. The parameter set includes all linear coefficients and variances, so that 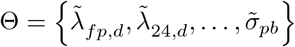. Taken together, the first term (Eq. S30) corresponds to the likelihood integrated over the latent factor *F*_*d*_ and *F*_*c*_, the second term (Eq. S31) the prior of the linear coefficients, and the third term (Eq. S32) the prior of the standard deviations.

The algebraic expression of the posteriors after integrating out the latent factors was obtained with Mathematica. Subsequently, the maximum a posteriori probability (MAP) parameters were obtained numerically by the gradient descent algorithms in Python library SciPy. Additionally, the covariance matrix of the MAP parameters was obtained by taking the inverse of the Hessian of the log-likelihood (i.e. **H**^*—1*^ where **H** = − ∇ ∇ ln *P* (Θ* **T**), and Θ* is the point estimate of the MAP parameter) [8]. The standard errors associated with the parameter estimates are the square root of the diagonal entries in this covariance matrix.

For each model, we generated a synthetic data set of 2,946 embryos, matching the size of the real cohort. Model parameters were fixed at their MAP values, while the latent factors and residual terms were drawn independently from a standard normal distribution. These values were plugged into the model equations to produce one complete timing vector per embryo. Variance and pair-wise correlation were then computed from the resulting 2,946 simulated timing vectors. This procedure is repeated 100 times to obtain the mean variance and correlation for all timings and timing pairs. These results are shown in Fig. S2 and Fig. 4 in the main text.

**Figure S2:**
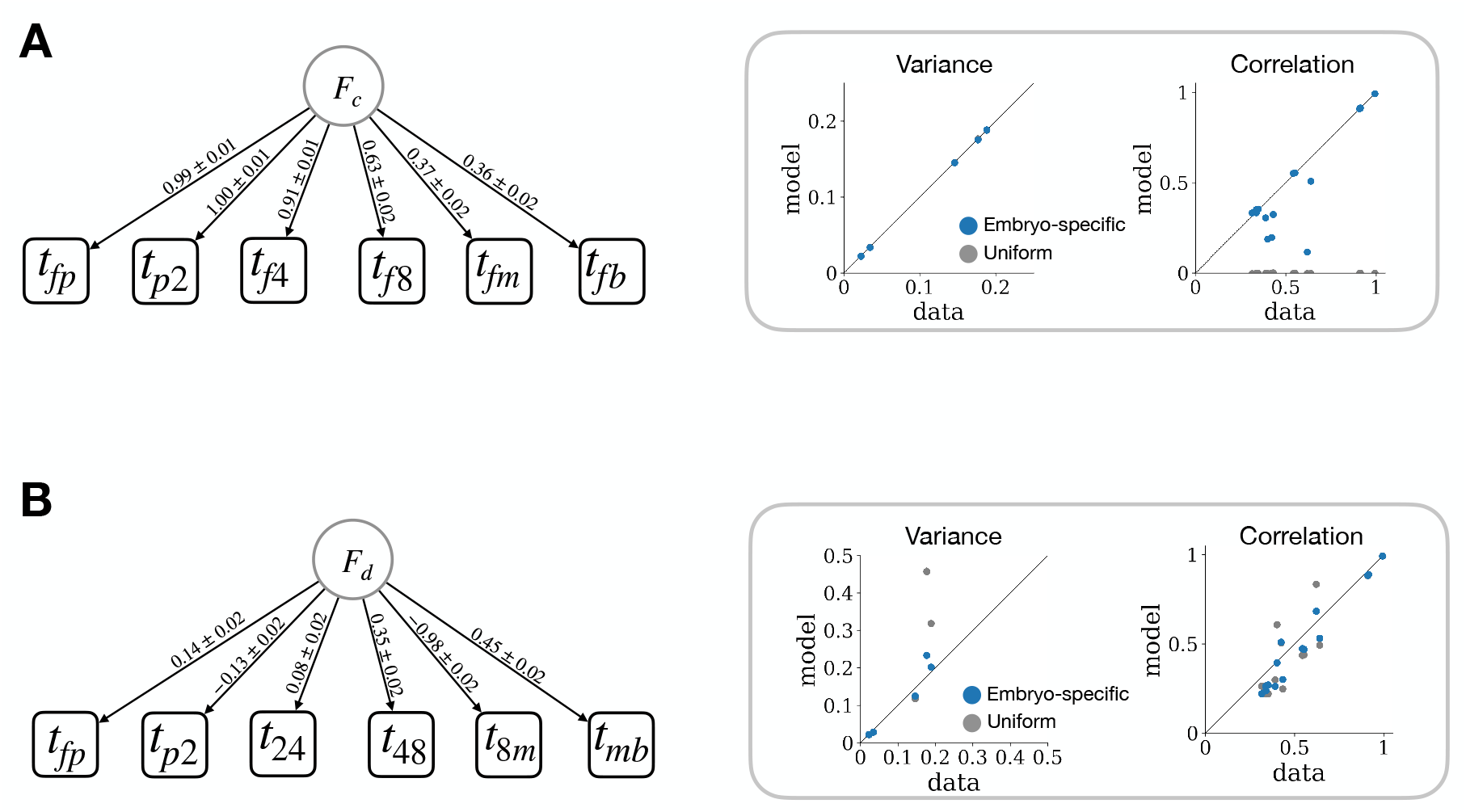
Maximum posterior fits for embryo-specific models. **A** Embryo-specific clock model. *B* Embryo-specific domino model. On the left of each panel, the numbers on the edges are the MAP estimates and the standard errors. On the right of each panel, the variance and correlation from data are plotted against that of the best-fit model. In the right panels, the uniform model values are shown in grey, and the corresponding embryo-specific model values are shown in blue.

#### 3.2 Model evidence

With MAP estimates for each model, we can also compare the goodness of fit between models as outlined in ref. [8]. The posterior probability of the model ℳ given data *T* is expressed as *P*(ℳ|**T**) ∝ *P*(**T**| ℳ) *P*(ℳ). Assuming that all models are equally likely to begin with, i.e. identical *P*(ℳ) for all ℳ, we compare the models by evaluating their evidence, *P*(**T**| ℳ). With Laplace approximation at the MAP estimate, *P*(**T**| ℳ) can be expressed as

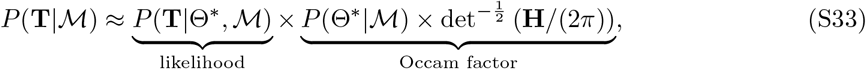

where **H** is the Hessian of the log-likelihood at the MAP estimate. As explained in ref. [8], the Occam factor quantifies how fine-tuned the parameters must be to achieve the best fit. More complex models with more parameters will result in better fits (i.e. higher likelihoods), but will also require more fine-tuning of the parameters (i.e.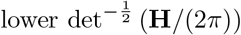)). The negative log model evidence of each model is presented in Table S2.

**Table S2:** Comparison of model evidence based on the Occam factor [8].

| Model name | Parameter count | Model evidence |
| --- | --- | --- |
| Uniform clock | 6 | 287 |
| Embryo-specific clock | 12 | 10017 |
| Uniform domino | 6 | 10778 |
| Embryo-specific domino | 12 | 11332 |

#### 3.3 Covariance-based tests

We evaluate the significance of the linear relationship between timing pairs in two ways: ordinary least squares (OLS) regression and bootstrapping.

OLS fit of variable *X* onto *Y* yields the linear slope that is equivalent to Cov(*X, Y*)*/*Var(*X*). Since Var(*X*) *>* 0, the slope is positive if and only if the covariance is positive. Similarly, the slope is positive if and only if the Pearson correlation, 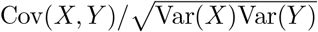, is positive. Thus, we use p-values from OLS regression to evaluate the significance of the covariance and correlation between timing pairs. Additionally, we set a lower threshold of 0.1 for significant correlations.

Although OLS is the standard method to quantify the significance of linear relationships, it depends on two key assumptions: (i) the data are independently sampled (ii) the residuals follow a normal distribution with fixed variance. Neither of these assumptions are guaranteed in the embryo timings. In particular, the embryos from the same patients tend to have similar rates of development (Fig. 5D and ref. [3]). Therefore, we also use bootstrapping to compute the significance of the correlations. Bootstrapping is conducted at the patient level with replacement *N*_*boot*_ = 10^*4*^ times. For the bootstrapped correlation coefficients, the measurement error was taken into account by following Eq. S2. The bootstrap p-value is defined as *p* = (min{*N*_*+*_, *N*_—_} + 1) */* (*N*_*boot*_) where *N*_*+*_ and *N*_—_ are the number of resamples in which the correlation difference was positive and negative, respectively. The plus one adjustment in the numerator prevents a p-value of 0 (Table S7).

**Table S3:** Significance of correlations shown in Fig. 2. In the second column, the p-values for the slope from linear regression are shown in parenthesis, after being corrected for multiple hypothesis testing with the Bonferroni method for *n* = 39 tests in this table. In the third column, the Bonferroni adjusted p-values in parenthesis are derived from bootstrapping. Significant results are set by *p*\* < 10^*—2*^ for both tests. The timing pairs with a non-significant linear relationship are colored grey.

|  | **Corr. coef.** | **Boot. corr. coef.** |
| --- | --- | --- |
| Clock model |  |  |
| $t_{fp}, t_{f2}$ | 0.99 (0.0e+00) | 1.00 (3.9e-03) |
| $t_{fp}, t_{f4}$ | 0.90 (0.0e+00) | 0.92 (3.9e-03) |
| $t_{f2}, t_{f4}$ | 0.91 (0.0e+00) | 0.91 (3.9e-03) |
| $t_{fp}, t_{f8}$ | 0.62 (5.1e-313) | 0.73 (3.9e-03) |
| $t_{f2}, t_{f8}$ | 0.63 (4.8e-319) | 0.74 (3.9e-03) |
| $t_{f4}, t_{f8}$ | 0.74 (0.0e+00) | 0.87 (3.9e-03) |
| $t_{fp}, t_{fm}$ | 0.36 (1.7e-87) | 0.38 (3.9e-03) |
| $t_{f2}, t_{fm}$ | 0.38 (1.8e-97) | 0.40 (3.9e-03) |
| $t_{f4}, t_{fm}$ | 0.45 (2.0e-148) | 0.48 (3.9e-03) |
| $t_{f8}, t_{fm}$ | 0.49 (6.1e-176) | 0.56 (3.9e-03) |
| $t_{fp}, t_{fb}$ | 0.34 (3.2e-81) | 0.37 (3.9e-03) |
| $t_{f2}, t_{fb}$ | 0.36 (1.4e-87) | 0.38 (3.9e-03) |
| $t_{f4}, t_{fb}$ | 0.46 (1.2e-153) | 0.48 (3.9e-03) |
| $t_{f8}, t_{fb}$ | 0.50 (2.0e-186) | 0.59 (3.9e-03) |
| $t_{fm}, t_{fb}$ | 0.69 (0.0e+00) | 0.76 (3.9e-03) |
| Domino model |  |  |
| $t_{p2}, t_{fp}$ | -0.08 (2.7e-04) | 0.04 (1.0e+00) |
| $t_{24}, t_{fp}$ | 0.25 (7.1e-42) | 0.29 (3.9e-03) |
| $t_{24}, t_{p2}$ | -0.04 (6.5e-01) | -0.17 (3.9e-03) |
| $t_{48}, t_{fp}$ | 0.16 (1.6e-17) | 0.24 (3.9e-03) |
| $t_{48}, t_{p2}$ | 0.02 (1.0e+00) | 0.17 (5.6e-02) |
| $t_{48}, t_{24}$ | 0.27 (2.7e-47) | 0.48 (3.9e-03) |
| $t_{8m}, t_{fp}$ | -0.14 (6.6e-13) | -0.17 (3.9e-03) |
| $t_{8m}, t_{p2}$ | 0.13 (1.2e-10) | 0.26 (3.9e-03) |
| $t_{8m}, t_{24}$ | -0.07 (2.7e-03) | -0.11 (3.9e-03) |
| $t_{8m}, t_{48}$ | -0.35 (2.5e-83) | -0.14 (1.5e-02) |
| $t_{mb}, t_{fp}$ | -0.02 (1.0e+00) | -0.01 (1.0e+00) |
| $t_{mb}, t_{p2}$ | -0.06 (4.6e-02) | -0.18 (3.9e-03) |
| $t_{mb}, t_{24}$ | 0.06 (3.2e-02) | 0.06 (3.6e-01) |
| $t_{mb}, t_{48}$ | 0.01 (1.0e+00) | 0.11 (8.9e-02) |
| $t_{mb}, t_{8m}$ | -0.45 (4.4e-142) | -0.43 (3.9e-03) |
| Miscellaneous |  |  |
| $t_{pm}, t_{fp}$ | 0.00 (1.0e+00) | 0.01 (1.0e+00) |
| $t_{pb}, t_{fp}$ | -0.01 (1.0e+00) | 0.00 (1.0e+00) |
| $t_{pm}, t_{p2}$ | 0.18 (1.1e-20) | 0.31 (3.9e-03) |
| $t_{2m}, t_{p2}$ | 0.13 (5.5e-11) | 0.28 (3.9e-03) |
| Continued on next page |  |  |

Table S3 – continued from previous page
|  | Corr. coef. | Boot. corr. coef. |
| --- | --- | --- |
| $t_{4m}, t_{p2}$ | 0.14 (1.2e-13) | 0.33 (3.9e-03) |
| $t_{pb}, t_{p2}$ | 0.13 (1.7e-10) | 0.17 (3.9e-03) |
| $t_{2b}, t_{p2}$ | 0.08 (6.7e-04) | 0.14 (2.2e-02) |
| $t_{4b}, t_{p2}$ | 0.09 (1.8e-05) | 0.18 (3.9e-03) |
| $t_{8b}, t_{p2}$ | 0.08 (8.7e-04) | 0.12 (2.1e-01) |
|  | Corr. coef. | Boot. corr. coef. |

#### 3.4 Partial covariance-based tests

We employ statistical tests that involve standard multiple linear regression and bootstrapping to evaluate the embryo-specific clock and domino models. Here, we provide an illustrative example to derive these statistical tests with the embryo-specific clock model timings (Eq. S9). Consider the equations for the times from fertilization to the 2-cell, 4-cell, and 8-cell,

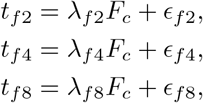

where the *λ* terms are the linear coefficients of the latent factor *F*_*c*_, and the *ϵ* terms represent mutually independent noise. Without loss of generality, we assume that the timings are normalized by their respective means and that the latent factor *F*_*c*_ is distributed with mean 0 and variance 1. Then, let *ϵ*_*f2*|*f4*_ and *ϵ*_*f8*|*f4*_ be the respective residuals from the linear fit of *t*_*f2*_ and *t*_*f8*_ by *t*_*f4*_, so that

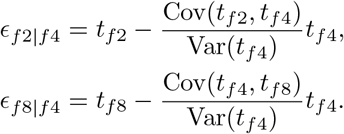

The covariance between *ϵ*_*f2*|*f4*_ and *ϵ*_*f8*|*f4*_, or the partial covariance between *t*_*f2*_ and *t*_*f8*_ given *t*_*f4*_, can then be expressed as

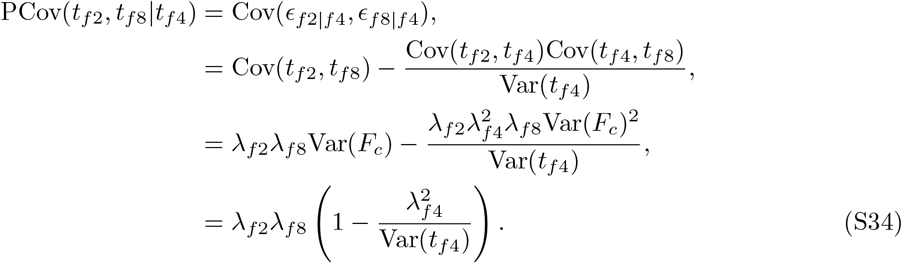

To obtain the last and second-to-last lines, we use the assumptions that *ϵ*_*f2*_, *ϵ*_*f4*_, *ϵ*_*f8*_ are all independent from each other and that Var(*F*_*c*_) = 1. By construction, 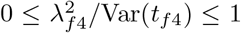. Thus, the covariance between residuals Cov(*ϵ*_*f8*|*f4*_, *ϵ*_*f2*|*f4*_) is positive if and only if Cov(*t*_*f2*_, *t*_*f8*_) = *λ*_*f2*_*λ*_*f8*_ is positive. We compute the partial covariance Cov(*r*_*f8*|*f4*_, *r*_*f2*|*f4*_) and the covariance Cov(*t*_*f2*_, *t*_*f8*_) by bootstrapping at the patient level. In all of the 10^*4*^ bootstrapped calculations, PCov(*t*_*f2*_, *t*_*f8*_|*t*_*f4*_) < 0 < Cov(*t*_*fp*_, *t*_*f8*_), thereby ruling out the embryo-specific clock model. Note that for this test, we did not assume any specific forms for the distribution of the residuals and the latent factor *F*_*c*_. All comparisons between the covariance and partial covariance in the clock and domino models are shown in Table S4. Also, note that the expression for the fraction variance explained, which is used to test the embryo-specific domino model in the main text, is obtained simply by dividing both sides of Eq. S34 by Cov(*t*_*f2*_, *t*_*f8*_). The bootstrapped values of the fraction of variance explained are shown in Table S5.

We can perform a similar analysis with multiple linear regression, by fitting *t*_*f8*_ as a function of *t*_*f2*_ and *t*_*f4*_. Here, note that the following equivalence relations hold: (i) the slope between *t*_*f2*_ and *t*_*f8*_ is positive if and only if Cov(*t*_*f2*_, *t*_*f8*_) *>* 0, and (ii) the partial coefficient between *t*_*f2*_ and *t*_*f8*_ given *t*_*f4*_ is positive if and only if PCov(*t*_*f2*_, *t*_*f8*_|*t*_*f4*_) *>* 0. Therefore, from Eq. S34, the partial coefficient between *t*_*f2*_ and *t*_*f8*_ given *t*_*f4*_ is positive if and only if the slope between *t*_*f2*_ and *t*_*f8*_ is positive, and the same holds for negative slopes. The significance of the slopes and partial coefficients evaluated by p-values from multiple linear regression are also shown in Table S4.

**Table S4:**
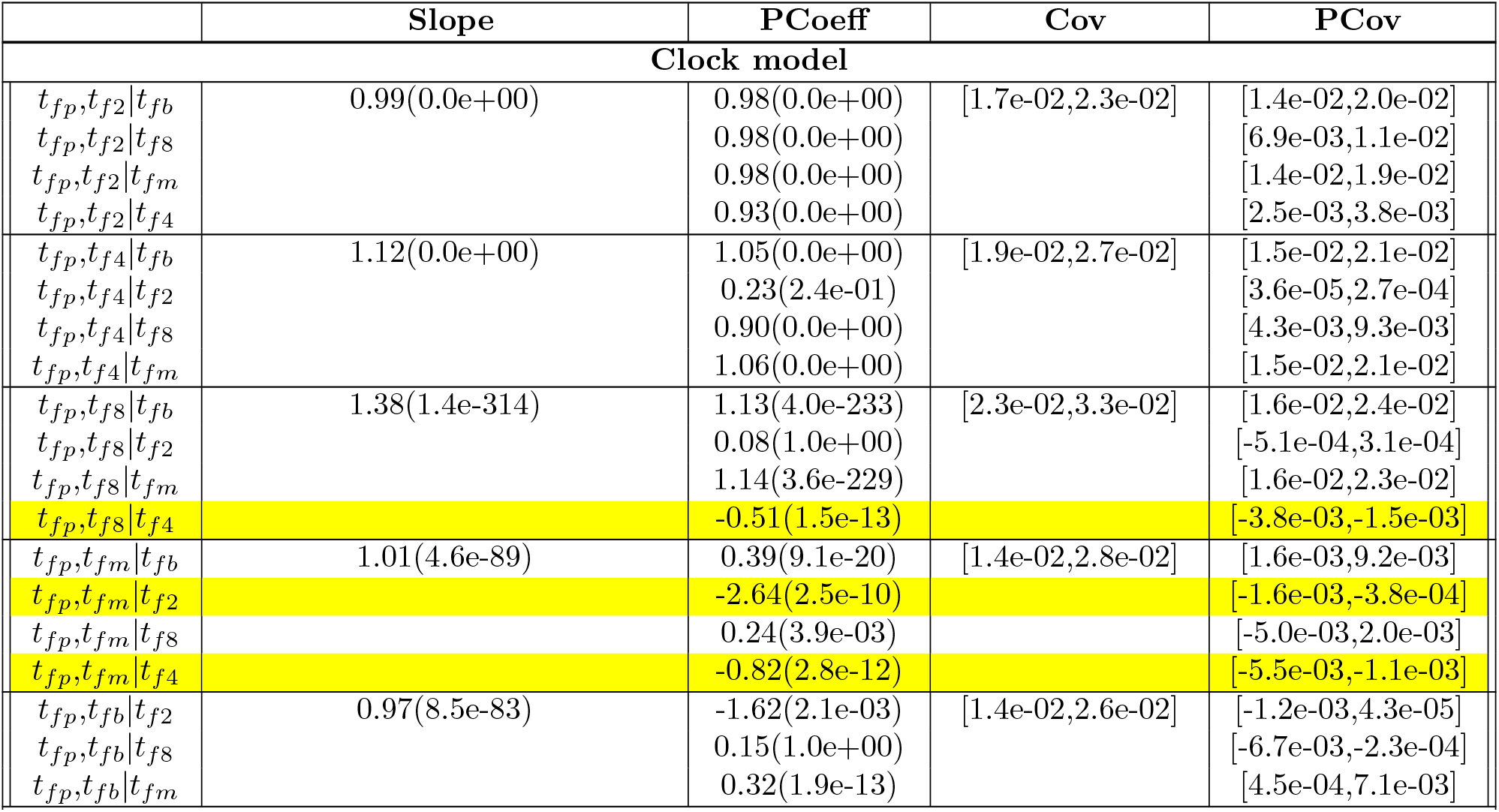
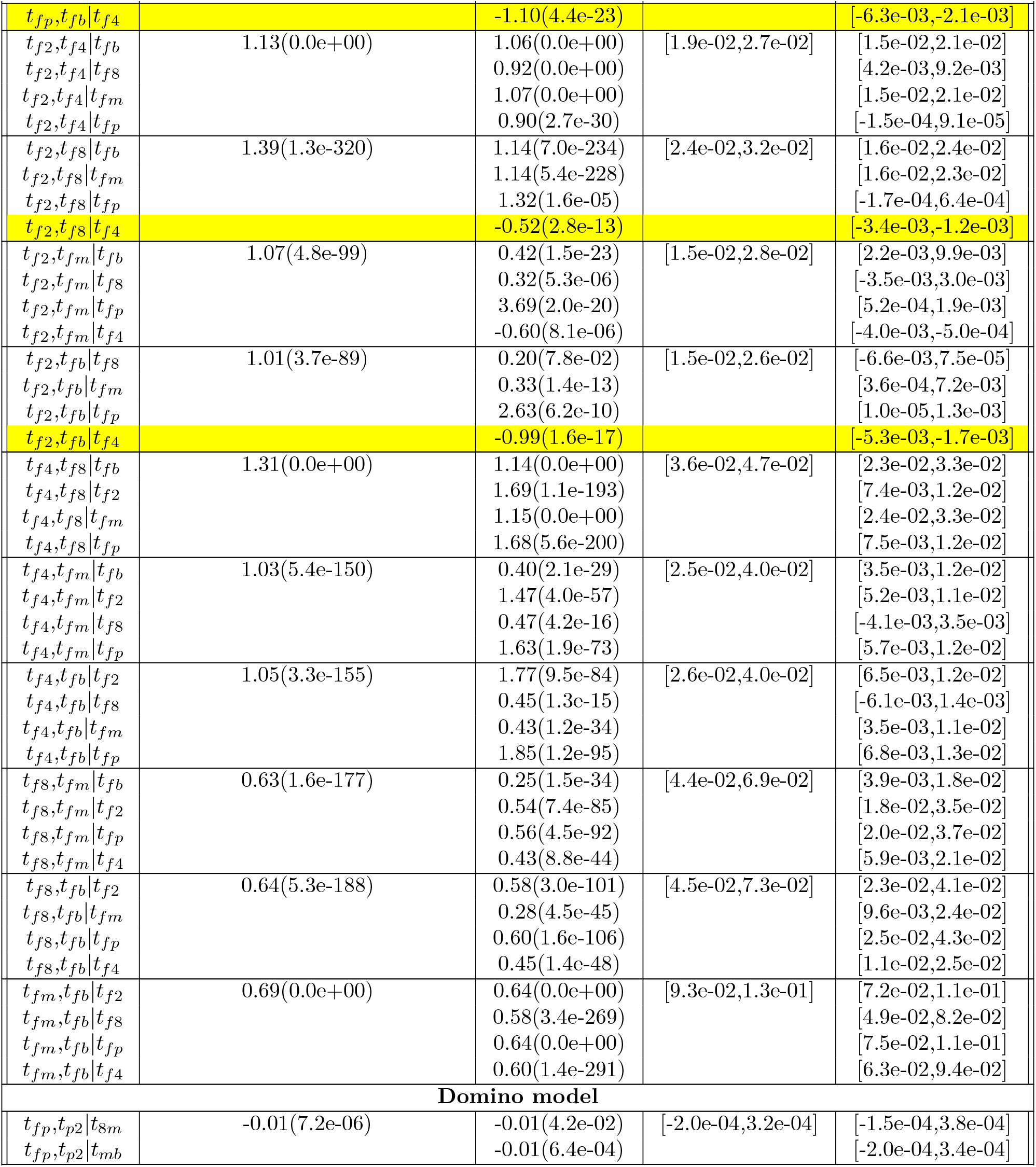
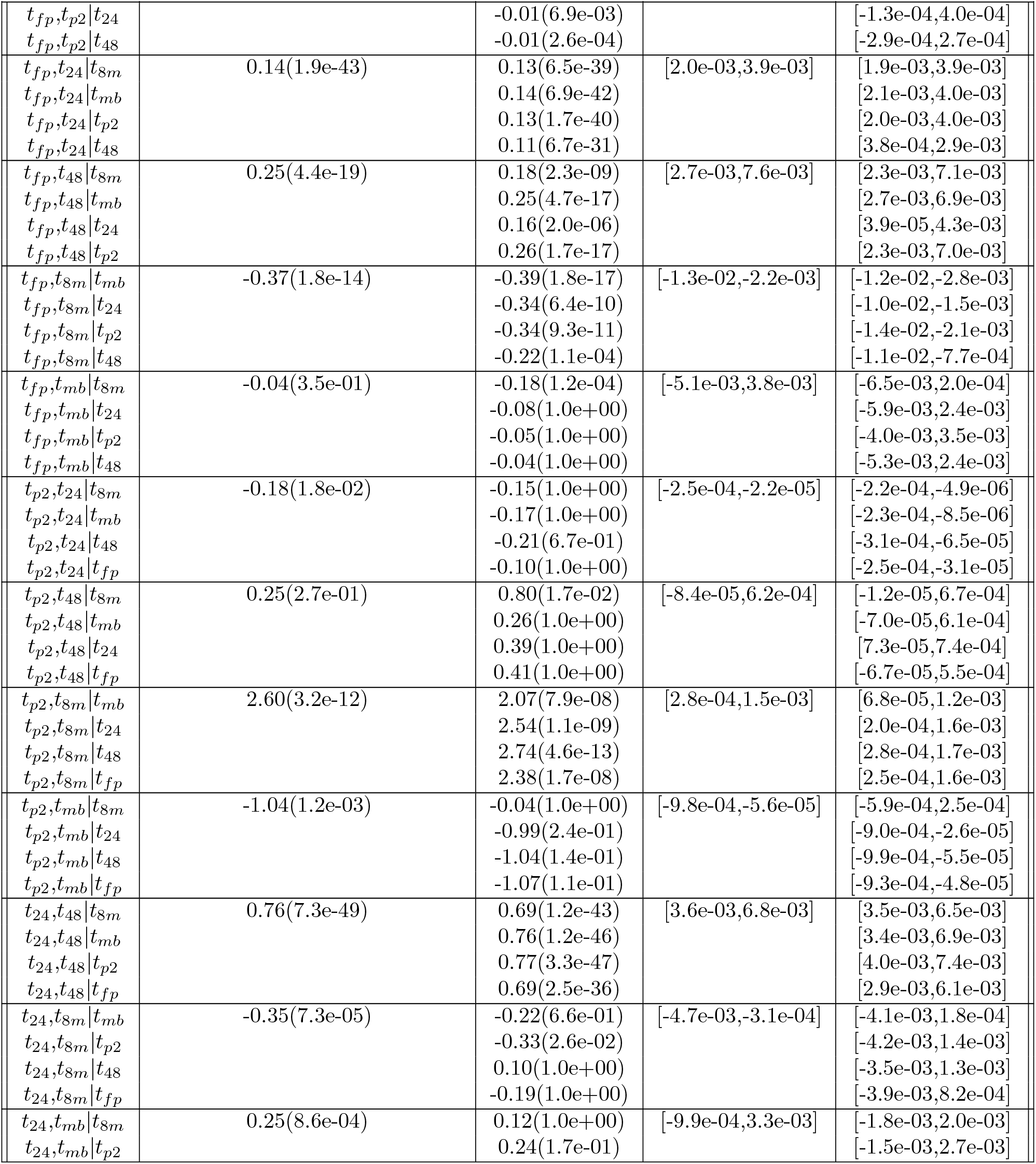
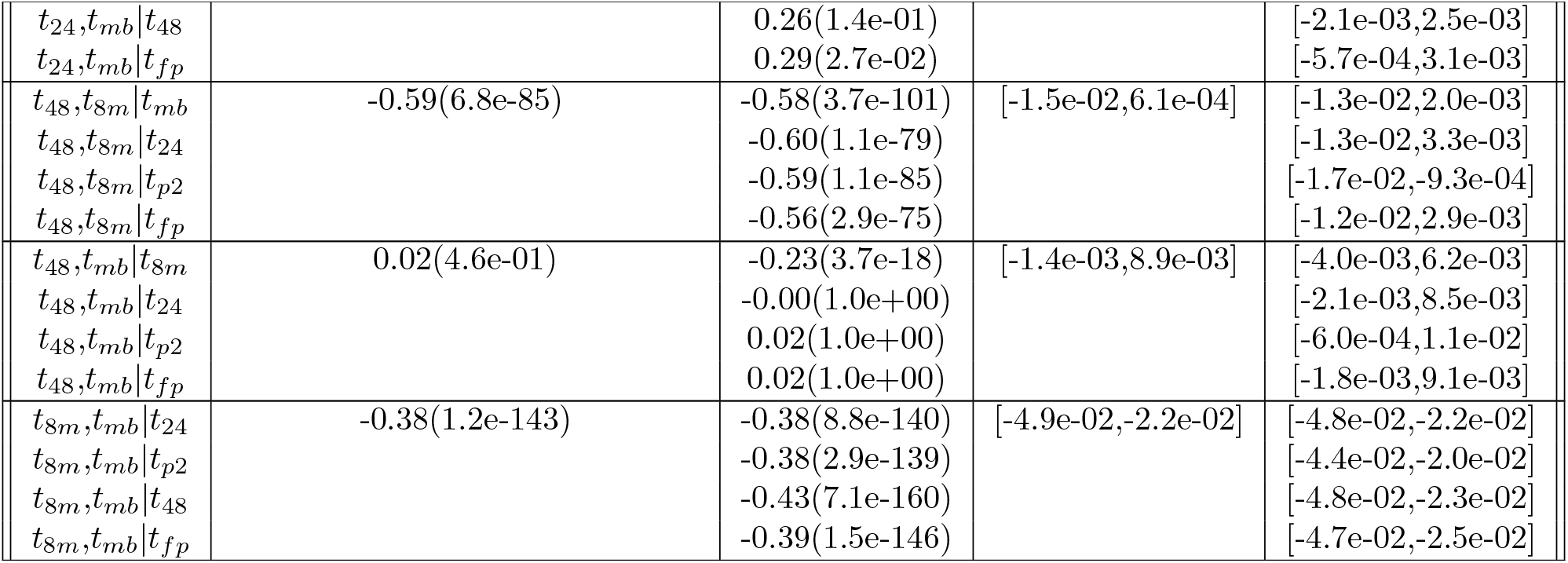
Marginal and partial covariance. () in the second column denotes the p-values for the pairwise slope without multiple hypothesis correction. () in the third column denotes the Bonferroni corrected p-values for *n* = 120 test for the partial coefficients obtained from multiple linear regression. The fourth and fifth columns denote the full range of covariance and partial co-variance values from bootstrapping. Instances where marginal and partial covariance have opposite signs with statistical significance are colored yellow, with significant results set by *p*\* < 10^*—2*^.

**Table S5:** Fraction of variance explained by the latent factor computed to test the embryo-specific domino model. The fraction of variance explained, computed as 1 — PCov(*t*_*X*_, *t*_*Y*_ |*t*_*Z*_)*/*Cov(*t*_*X*_, *t*_*Y*_), is shown only when the correlation between the pair (*t*_*X*_, *t*_*Y*_) is significant. The full range of values from bootstrapping at the patient level 10^*4*^ times is shown. Instances where the bootstrapped range do not overlap are highlighted in yellow

|  | Fraction of variance explained |
| --- | --- |
| $t_{48}, t_{24} \vert t_{fp}$ | [6.5e-02, 2.6e-01] |
| $t_{p2}, t_{8m} \vert t_{fp}$ | [-2.1e-01, 1.0e-01] |
| $t_{mb}, t_{8m} \vert t_{fp}$ | [-4.4e-02, 5.4e-02] |
| $t_{fp}, t_{48} \vert t_{p2}$ | [-1.2e-01, 2.4e-01] |
| $t_{48}, t_{24} \vert t_{p2}$ | [-1.7e-01, 3.2e-02] |
| $t_{fp}, t_{8m} \vert t_{p2}$ | [-5.7e-01, 3.0e-01] |
| $t_{mb}, t_{8m} \vert t_{p2}$ | [3.5e-03, 3.1e-01] |
| $t_{fp}, t_{48} \vert t_{24}$ | [3.3e-01, 1.0e-00] |
| $t_{p2}, t_{8m} \vert t_{24}$ | [4.7e-03, 2.8e-01] |
| $t_{fp}, t_{8m} \vert t_{24}$ | [1.4e-02, 7.3e-01] |
| $t_{mb}, t_{8m} \vert t_{24}$ | [-6.9e-03, 6.7e-02] |
| $t_{p2}, t_{8m} \vert t_{48}$ | [-6.0e-01, 2.1e-02] |
| $t_{fp}, t_{8m} \vert t_{48}$ | [-1.4e-01, 5.2e-01] |
| $t_{mb}, t_{8m} \vert t_{48}$ | [-7.4e-03, 1.1e-01] |
| $t_{fp}, t_{24} \vert t_{48}$ | [1.7e-01, 7.4e-01] |
| $t_{48}, t_{fp} \vert t_{8m}$ | [-2.2e-02, 3.5e-01] |
| $t_{48}, t_{24} \vert t_{8m}$ | [-3.1e-03, 1.1e-01] |
| $t_{fp}, t_{24} \vert t_{8m}$ | [3.8e-03, 1.6e-01] |
| $t_{48}, t_{fp} \vert t_{mb}$ | [-8.1e-02, 6.1e-02] |
| Continued on next page |  |

Table S5 – continued from previous page
|  | Fraction of variance explained |
| --- | --- |
| $t_{48}, t_{24} \vert t_{mb}$ | $[-8.3e-03, 5.6e-02]$ |
| $t_{p2}, t_{8m} \vert t_{mb}$ | $[3.7e-02, 7.7e-01]$ |
| $t_{8m}, t_{fp} \vert t_{mb}$ | $[-5.3e-01, 2.1e-01]$ |
| $t_{fp}, t_{24} \vert t_{mb}$ | $[-3.7e-02, 3.2e-02]$ |
|  | Fraction of variance explained |

**Figure S3:**
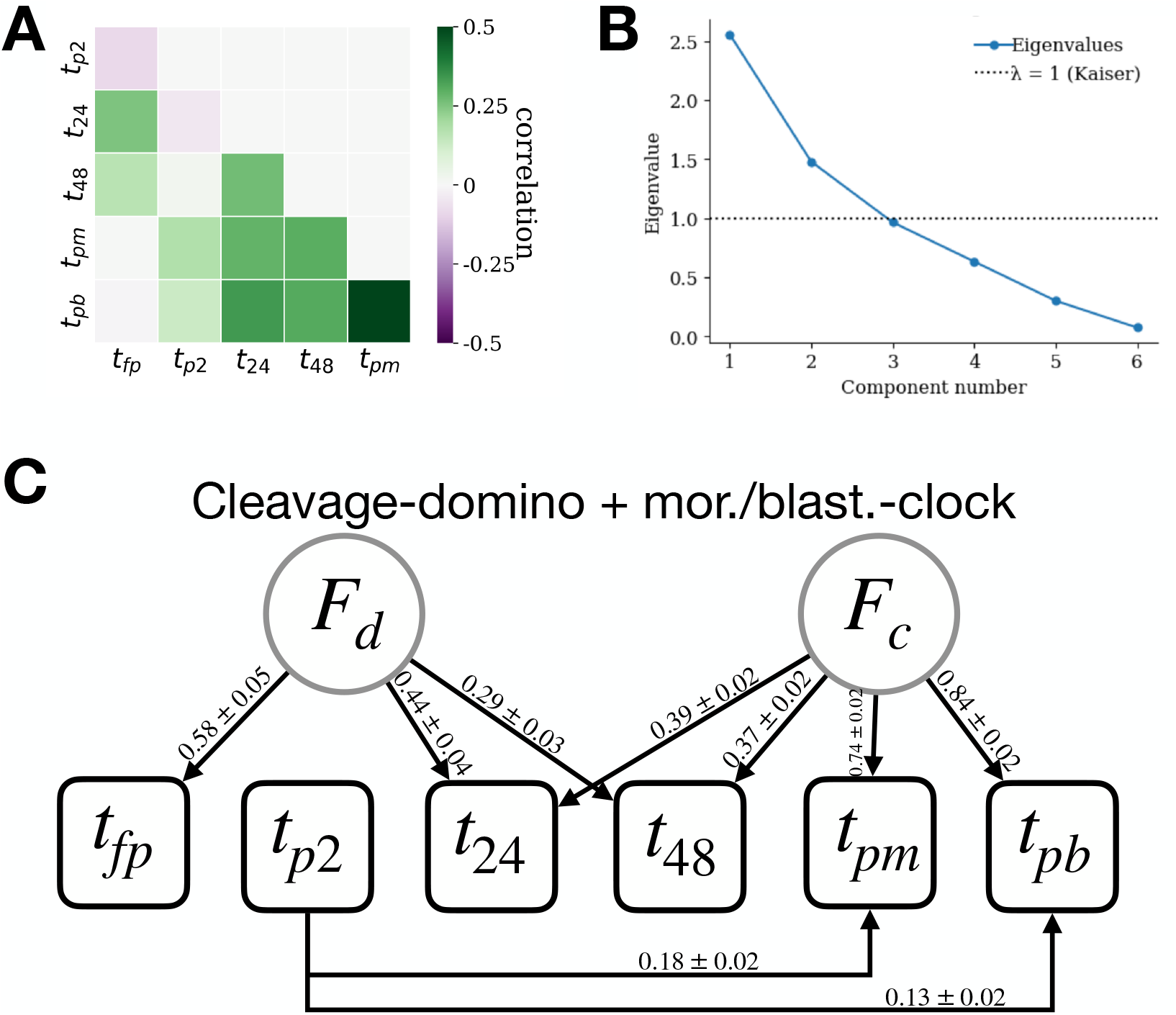
Cleavage-domino+mor./blast.-clock model. **A** The correlation matrix of the timings. **B** The eigenvalues of the correlation matrix, showing that only two principal components have eigenvalues larger than 1 (i.e. the Kaiser criteria). **C** Numbers on the edges are the maximum a posteriori probability (MAP) estimates of the corresponding parameters. denotes the standard errors.

**Table S6:**
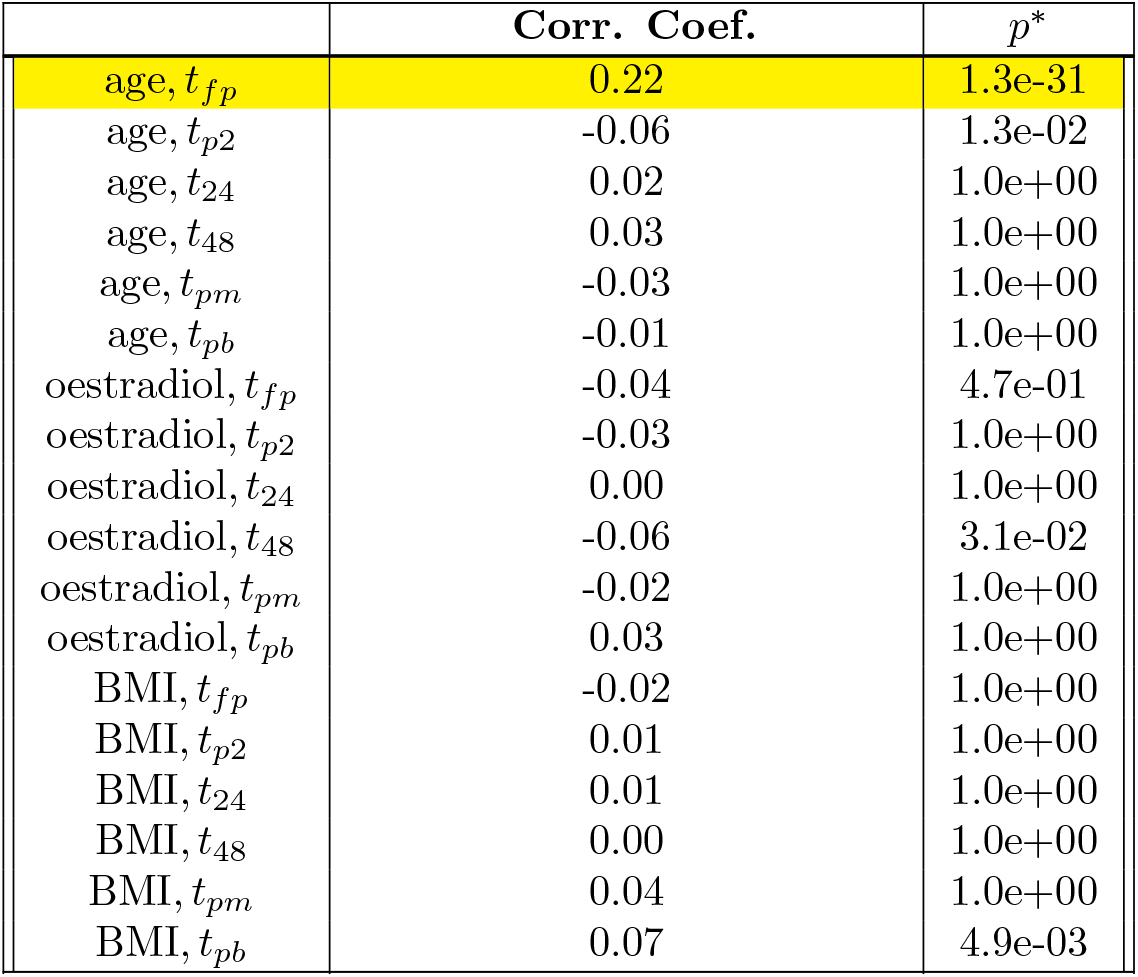
Correlation between external variables and timings. *p*\* denotes Bonferroni correction adjusted p-values for *n* = 18 tests in this table. Statistically significant correlations (*p*\* < 10^*—2*^) are marked in yellow. For age, there are 2946 embryos included in the calculations. For oestradiol and BMI, there were 2920 and 2626 embryos, respectively.

#### 3.5 Fraction of variance explained by patient and treatment cycle factors

We provide detail on how the fraction of variance explained by patient and treatment cycle factors are obtained from Bayesian hierarchical regression, by working through an illustrative example with *t*_*fp*_. Results for all other developmental stages are computed identically.

We denote the total number of patients by *M*, with each patient *k* undergoing *N*^*k*^ treatment cycles for *k* = 1, …, *M*. Each treatment cycle *j* for patient *k* includes *n*^*kj*^ embryos, for *j* = 1, …, *N*_*k*_. Without loss of generality, we normalize the timings by the mean and standard deviation throughout the embryo population, so that the global average timing is set to *µ*^*g*^ = 0. For each patient *k*, we define their mean timing as *µ*^*k*^, and for each treatment cycle *j* within patient *k*, we define the cycle-specific mean as *µ*^*kj*^. The normalized timing of embryo *i* in cycle *j* of patient *k* is denoted as 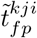. Then, we assume a hierarchical model with the following nested structure:

- Patient mean: 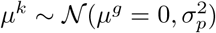
- Cycle mean: 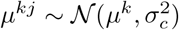
- Embryo value: 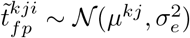

In this model, the total variance in embryo timing is decomposed as:

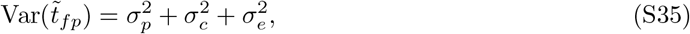

where 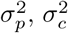, and 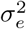 denote the variance associated with the patient, treatment cycle, and embryo, respectively. We define Θ to be the set of all model parameters: 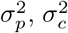, and 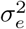, as well as the latent means *µ*^*k*^ and *µ*^*kj*^ for all patients and cycles. Then, the posterior distribution of the full parameter set Θ given the observed embryo timing data **T** is proportional to

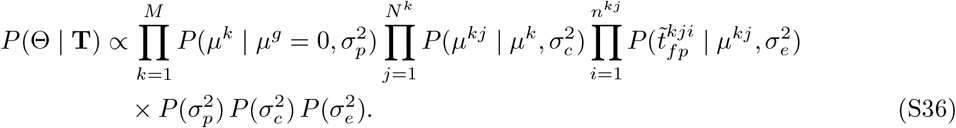

The right hand side of the first line correspond to likelihoods at each level of the hierarchy. The terms on the second line correspond to the priors for 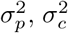, and 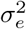, which are chosen to be uninformative through the uniform distributions with support [0.001, 10]. Markov Chain Monte Carlo (MCMC) is performed to produce the maximum a posteriori (MAP) estimates and associated credible intervals. We used 14000 MCMC steps, after discarding the first 4000 steps as burn-in. **T** includes only treatment cycles that contain more than one embryo, and patients that underwent more than one treatment cycle, ensuring that the variance estimates are identifiable at each level. The dataset for this analysis includes 2,454 embryos from 686 treatment cycles across 520 patients. Results are summarized in Fig. 5D.

#### 3.6 Significance of the association between timings and clinical pregnancy outcome

We employ a linear model to evaluate the association between developmental timing and clinical pregnancy outcome, from the selection of transferred embryos as described in supplementary section 1.4. The timings are all normalized by their respective mean and standard deviation. For patient *k* and embryo *i*, the normalized timings are denoted by 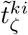 for developmental stage ζ = *f p, p*2, 24, 48, *pm, pb*. We fit the OLS model

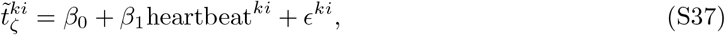

where heartbeat^*ki*^ equals 1 for a successful clinical pregnancy outcome, and 0 otherwise. There may be correlations among the developmental timings and the clinical pregnancy outcomes from the same patient. Consequently, the conventional standard error for the maximum-likelihood estimate 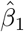 is inappropriate. We therefore obtain a cluster-robust standard error for 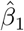 using the Liang–Zeger estimator [7], treating each patient as one cluster. The resulting test statistic

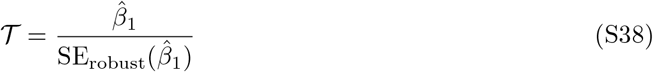

is compared with a Student-t distribution with *C* degrees of freedom, where *C* is the number of patients. The resulting two-sided p-value for the null hypothesis *β*_*1*_ = 0 is shown in Table S7.

To further corroborate this result, we perform patient-level bootstrapping. We draw patients with replacement *N*_*boot*_ = 10^*4*^ times, recompute the mean difference in the timing variable between the embryos that resulted in clinical pregnancy and those that did not, and record its sign. The bootstrap p-value is defined as *p*_*boot*_ = (min *N*_*+*_, *N*_—_ + 1) */* (*N*_*boot*_), where *N*_*+*_ and *N*_—_ are the number of bootstrapped samples in which the mean difference was positive and negative, respectively. The plus one adjustment in the denominator prevents a p-value of 0 (Table S7).

**Table S7:** Association between clinical pregnancy rate and developmental timing. The number of embryos included for each calculation is shown in (). *p*\* denotes Bonferroni correction adjusted p-values for *n* = 24 tests in this table. 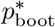 denotes the p-values from bootstrapping after multiple hypothesis correction for *n* = 24 tests. Timings with statistically significant differences between clinical pregnancy outcomes evaluated by both metrics, i.e. 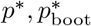, are marked in yellow.

| timing | fetal heartbeat | no fetal heartbeat | $p^*$ | $p_{boot}^*$ |
| --- | --- | --- | --- | --- |
| $t_{fp}$ | 9.79e-01 (184) | 1.02e+00 (2001) | 6.01e-04 | 2.40e-03 |
| $\mu_{fp}^{pat}$ | 9.91e-01 (156) | 1.03e+00 (1685) | 4.72e-02 | 1.44e-02 |
| $\delta t_{fp}$ | -1.64e-02 (156) | -1.11e-02 (1685) | 1.00e+00 | 1.00e+00 |
| $\mu_{fp}^{sib}$ | 9.96e-01 (140) | 1.04e+00 (1190) | 1.86e-01 | 1.78e-01 |
| $t_{p2}$ | 1.11e-01 (184) | 1.13e-01 (1990) | 1.00e+00 | 1.00e+00 |
| $\mu_{p2}^{pat}$ | 1.12e-01 (154) | 1.13e-01 (1630) | 1.00e+00 | 1.00e+00 |
| $\delta t_{p2}$ | -1.54e-03 (154) | -7.62e-04 (1630) | 1.00e+00 | 1.00e+00 |
| $\mu_{p2}^{sib}$ | 1.13e-01 (138) | 1.15e-01 (1077) | 1.00e+00 | 1.00e+00 |
| $t_{24}$ | 5.10e-01 (183) | 5.32e-01 (1943) | 1.70e-04 | 2.40e-03 |
| $\mu_{24}^{pat}$ | 5.16e-01 (155) | 5.40e-01 (1581) | 1.94e-04 | 2.40e-03 |
| $\delta t_{24}$ | -8.56e-03 (155) | -9.74e-03 (1581) | 1.00e+00 | 1.00e+00 |
| $\mu_{24}^{sib}$ | 5.14e-01 (141) | 5.56e-01 (1033) | 1.57e-07 | 2.40e-03 |
| $t_{48}$ | 7.11e-01 (156) | 7.46e-01 (1446) | 6.20e-01 | 3.94e-01 |
| $\mu_{48}^{pat}$ | 7.45e-01 (135) | 7.72e-01 (1165) | 1.00e+00 | 1.00e+00 |
| $\delta t_{48}$ | -4.85e-02 (135) | -3.04e-02 (1165) | 1.00e+00 | 1.00e+00 |
| $\mu_{48}^{sib}$ | 7.75e-01 (125) | 8.22e-01 (816) | 2.98e-01 | 8.66e-01 |
| $t_{pm}$ | 2.38e+00 (80) | 2.45e+00 (410) | 1.00e+00 | 1.00e+00 |
| $\mu_{pm}^{pat}$ | 2.51e+00 (67) | 2.54e+00 (316) | 1.00e+00 | 1.00e+00 |
| $\delta t_{pm}$ | -9.89e-02 (67) | -6.44e-02 (316) | 1.00e+00 | 1.00e+00 |
| $\mu_{pm}^{sib}$ | 2.57e+00 (67) | 2.57e+00 (277) | 1.00e+00 | 1.00e+00 |
| $t_{pb}$ | 3.06e+00 (64) | 3.13e+00 (234) | 1.00e+00 | 1.00e+00 |
| $\mu_{pb}^{pat}$ | 3.20e+00 (54) | 3.21e+00 (170) | 1.00e+00 | 1.00e+00 |
| $\delta t_{pb}$ | -1.05e-01 (54) | -7.67e-02 (170) | 1.00e+00 | 1.00e+00 |
| $\mu_{pb}^{sib}$ | 3.26e+00 (54) | 3.29e+00 (162) | 1.00e+00 | 1.00e+00 |

